# Adaptation of codon and amino acid use for translational functions in highly expressed cricket genes

**DOI:** 10.1101/2020.08.31.276477

**Authors:** Carrie A. Whittle, Arpita Kulkarni, Nina Chung, Cassandra G. Extavour

**Author notes:** Corresponding author: Cassandra G. Extavour.

## Abstract

**Background:** For multicellular organisms, much remains unknown about the dynamics of synonymous codon and amino acid use in highly expressed genes, including whether their use varies with expression in different tissue types and sexes. Moreover, specific codons and amino acids may have translational functions in highly transcribed genes, that largely depend on their relationships to tRNA gene copies in the genome. However, these relationships and putative functions are poorly understood, particularly in multicellular systems.

**Results:** Here, we rigorously studied codon and amino acid use in highly expressed genes from reproductive and nervous system tissues (male and female gonad, somatic reproductive system, brain, ventral nerve cord, and male accessory glands) in the cricket *Gryllus bimaculatus*. We report an optimal codon, defined as the codon preferentially used in highly expressed genes, for each of the 18 amino acids with synonymous codons in this organism. The optimal codons were largely shaped by selection, and their identities were mostly shared among tissue types and both sexes. However, the frequency of optimal codons was highest in gonadal genes. Concordant with translational selection, a majority of the optimal codons had abundant matching tRNA gene copies in the genome, but sometimes obligately required wobble tRNAs. We suggest the latter may comprise a mechanism for slowing translation of abundant transcripts, particularly for cell-cycle genes. Non-optimal codons, defined as those least commonly used in highly transcribed genes, intriguingly often had abundant tRNAs, and had elevated use in a subset of genes with specialized functions (gametic and apoptosis genes), suggesting their use promotes the upregulation of particular mRNAs. In terms of amino acids, we found evidence suggesting that amino acid frequency, tRNA gene copy number, and amino acid biosynthetic costs (size/complexity) had all interdependently evolved in this insect model, potentially for translational optimization.

**Conclusions:** Collectively, the results strongly suggest that codon use in highly expressed genes, including optimal, wobble, and non-optimal codons, and their tRNAs abundances, as well as amino acid use, have been adapted for various functional roles in translation within this cricket. The effects of expression in different tissue types and the two sexes are discussed.

## Background

Synonymous codons in protein-coding genes are not used randomly [1]. The preferential use of synonymous codons per amino acid in highly transcribed genes, often called optimal codons, has been observed in diverse organisms including bacteria, fungi, plants and animals [2-14], including insects such as flies, mosquitoes, beetles and crickets [10, 11, 13, 15, 16]. When optimal codons co-occur with a high count of iso-accepting tRNA gene copies in the genome, which reflects an organism’s tRNA abundance [3-5, 12, 17-20], it suggests a history of selection favoring translational optimization [1, 3, 5, 12, 13, 20-25]. In multicellular organisms, unlike unicellular systems, genes can be expressed at different levels among tissue types and between the two sexes [16, 26-29]. Thus, in these organisms, codon use may be more complex, given that it is plausible that optimal codons may depend on the tissue type or sex in which a gene is expressed [11, 16, 22, 30, 31], and codon use could feasibly adapt to local tissue-dependent tRNA populations [30, 32, 33]. However, only minimal data are currently available about whether and how codon use varies with high expression in different tissue types and between the two sexes in multicellular organisms.

The limited data that are available suggest that codon use varies among genes transcribed in different tissues. We recently found, for example, that some optimal codons of highly transcribed genes differed among males and females for the testis, ovaries, gonadectomized-males and gonadectomized females, which may suggest adaptation of codon use to local tRNA populations in the beetle *T. casteneum* [16]. In addition, a study in *Drosophila melanogaster* showed that certain codons were preferentially used in the testis (CAG (Gln), AAG (Lys), CCC (Pro), and CGU (Arg)) as compared to other tissues such as the midgut, ovaries, and salivary glands, a result that was taken as support for the existence of tissue-specific tRNA populations [32] (see also an analysis of codon bias by [31]). Similar patterns of tissue-related use of specific codons have been inferred in humans [33, 34] and the plants *Arabidopsis thaliana* and *Oryza sativa* [30, 35]. Given the limited scope of organisms studied to date, however, further research is needed to determine whether the codon use varies among tissues across a broader scale of organisms. Tissues that are of particular importance for research include the gonads, which are key to reproductive success, and the brain, wherein the transcribed genes are apt to regulate male and female sexual behaviors [36-38]. Translational optimization of highly transcribed genes in these tissues may be particularly significant for an organism’s fitness.

While much of the focus on codon use in an organism’s highly expressed genes to date has centered on optimal codons [3, 5, 7, 12, 13, 16, 21-25, 39-41], and whether they have abundant matching tRNAs that may improve translation [3, 12, 13, 20-24], growing evidence suggests that other, less well studied, types of codon statuses could also play important translational roles [42-44]. In particular, even for codons that are not optimal, the supply-demand relationship between codons and tRNA abundances may regulate translation rates, possibly affecting protein functionality and abundance [42, 45-47]. For example, *in vivo* experimental research has shown that genes using codons requiring wobble tRNAs, which imprecisely match a codon at the third nucleotide site, exhibit slowed movement of ribosomes along mRNAs [42, 48, 49]. Similarly, non-optimal codons, defined as those codons that are least commonly used in highly transcribed genes (or sometimes defined as “rare” codons), particularly those non-optimal codons with few or no tRNAs in the cellular tRNA pool [16], may decelerate translation and thereby prevent ribosomal jamming [19] and also allow proper co-translational protein folding [44, 50-53]. In this regard, wobble codons, and non-optimal codons with few matching tRNA gene copies in the genome, may have significant translational roles, including roles in slowing translation.

In contrast to non-optimal codons that have few tRNAs, some evidence has emerged suggesting non-optimal codons may sometimes have abundant tRNAs, a relationship that may act to improve translation of specific gene mRNAs [16, 45]. For instance, in yeast (*Saccharomyces cerevisiae*), rare genomic codons exhibit enhanced use in stress genes, and tRNAs matching these codons have been found to be upregulated in response to stressful conditions, allowing improvement of their translation levels without any change in transcription rates [45]. In the red flour beetle, we recently reported that some non-optimal codons have abundant matching tRNA genes in the genome [16], and these codons are concentrated in a subset of highly transcribed genes with specific, non-random biological functions (e.g., olfactory or stress roles), which may together allow preferential translation of mRNAs of those particular genes [16]. Accordingly, given these findings, further studies of codon use patterns in highly expressed genes of multicellular organisms should expand beyond the focus on optimal codons *per se* [2, 3, 7-9, 12, 21, 39, 41], and explore the use and possible translational functions of non-optimal codons, distinguishing between those that have few and plentiful tRNAs, as well as the use of wobble codons [16].

While the investigation of amino acid use remains uncommon in multicellular organisms, the available sporadic studies suggest an association between amino acid use and gene expression level [21, 54, 55]. In insects, for example, an assessment of the biosynthetic costs of amino acid synthesis (size/complexity score for each of 20 amino acids as quantified by Dufton [56]) has shown that those amino acids with low costs tend to be more commonly used in genes with high transcription levels in the beetle *T. castaneum* [21]. Further, genome-wide studies in other arthropod models such as spiders (*Parasteatoda tepidariorum*) [55], and the study of partial available transcriptomes from milkweed bugs (*Oncopeltus fasciatus*), an amphipod crustacean (*Parhyale hawaiensis*) and crickets (*Gryllus bimaculatus*, using a single ovary/embryo dataset in this system) [10], were suggestive of the hypothesis that evolution may have typically favored a balance between minimizing the amino acid costs for production of abundant proteins with the need for certain (moderate cost) amino acids to ensure proper protein function (protein stability and/or functionality) [54]. Moreover, it has been found that amino acid use is correlated to their tRNA gene copy numbers in beetles [21], and in some other eukaryotes [17], a relationship that may be stronger in highly transcribed genes [17]. Thus, these various patterns raise the possibility of adaptation of amino acid use for translational optimization in multicellular organisms [17, 21, 55]. At present, further data is needed on amino acid use in highly expressed genes in multicellular systems, that include consideration of tRNA gene number, biosynthetic costs, and expression in different tissue types.

An emerging model system that provides opportunities for further deciphering the relationships between gene expression and codon and amino acid use is the two-spotted cricket *Gryllus bimaculatus*. Within insects, *Gryllus* is a hemimetabolous genus (Order Orthoptera) and has highly diverged from the widely studied model insect genus *Drosophila* (Order Diptera) [57, 58]. *G. bimaculatus* comprises a model for investigations in genetics [59, 60], germ line formation and development [61-63] and for molecular evolutionary biology [10, 64]. In the present study, we rigorously assess codon and amino acid use in highly transcribed genes of *G. bimaculatus* using its recently available annotated genome [65] and large-scale RNA-seq data from tissues of the male and female reproductive and nervous systems [64]. From our analyses, we demonstrate that optimal codons, those preferentially used in highly expressed genes, occur in this organism, are largely shaped by selection pressures, and are nearly identical across tissues. Based on analyses of codon and tRNA gene copy relationships, we find that a majority of optimal codons have abundant tRNAs, which is consistent with translational optimization in this species. However, some optimal codons obligately require the use of wobble tRNAs, which may act to slow translation, including for cell-cycle genes. Moreover, non-optimal codons, those codons rarely used in highly expressed genes, rather than usually having few tRNAs, often have abundant tRNAs, and thus may provide a system to upregulate the translation of specific mRNAs (for example, apoptosis gonadal genes), as has been proposed in yeast and beetles [16, 45]. Finally, with respect to amino acids, we find evidence to suggest that amino acid frequency, tRNA gene copy number, and amino acid biosynthetic costs have all interdependently evolved in this taxon, possibly for translational optimization.

## Results and Discussion

For our study, codon and amino acid use in *G. bimaculatus* was assessed using genes from its recently available annotated genome [65]. We included all 15,539 *G. bimaculatus* protein-coding genes (CDS, longest CDS per gene) that had a start codon and were >150bp. Gene expression was assessed using RNA-seq data from four adult male and female tissue types, the gonad (testis for males, ovaries for females), somatic reproductive system (for males this includes the pooled vasa deferentia, seminal vesicle and ejaculatory duct and for females includes the spermathecae, common oviduct, and bursa), brain and ventral nerve cord (Additional file 1: Table S1; [64]). The male accessory glands were included for study, but were separated from the other male reproductive system to prevent overwhelming, or skewing, the types of transcripts detected in the former tissues [64]. The trimmed reads in Additional file 1: Table S1 were mapped to the 15,539 annotated G. *bimaculatus* genes independently for each of the nine tissue types under study and the expression level, or FPKM, was determined per gene.

To identify the optimal and non-optimal codons in *G. bimaculatus*, we compared codon use in highly versus lowly expressed genes [2, 7, 9, 10, 15, 16, 39, 66, 67]. For each CDS, the relative synonymous codon usage (RSCU) was determined for all codons for each amino acid with synonymous codons, whereby RSCU values >1 and <1 respectively indicate greater and lower use of a synonymous codon than that expected under equal codon use, and elevated values of codons for each amino acid indicate more frequent usage [18]. The ΔRSCU=RSCU_Mean Highly Expressed CDS_-RSCU_Mean Low Expressed CDS_ was used to define the primary optimal codon as the codon with the largest positive and statistically significant ΔRSCU value per amino acid [2, 7, 9, 10, 15, 16, 39]. The primary non-optimal codon was defined as the codon with the largest negative and statistically significant ΔRSCU value per amino acid [16]. In the following sections, we first thoroughly describe the optimal codons identified in this cricket species, including an assessment of the variation in expression among tissue types, and the role of selection versus mutation in shaping the optimal codons. Subsequently, we thoroughly evaluate the relationships between optimal codons and the non-optimal codons and their matching tRNA gene counts in the genome to ascertain plausible functional roles. We then consider the amino acid use and tRNA relationships in highly expressed genes of this taxon.

### Optimal Codons are Shared Across the Nine Distinct Tissues in *G. bimaculatus*

The organism-wide optimal codons were identified for *G. bimaculatus* using ΔRSCU for genes with the top 5% average expression levels across all nine studied tissues (cutoff was 556.2 FPKM) versus the 5% of genes with the lowest average expression levels (among all 15,539 genes under study) and are shown in Table 1. Based on ΔRSCU we report a primary optimal codon for all of the 18 amino acids with synonymous codons, each of which ended at the third position in an A (A3) or T (T3) nucleotide (boldface and underlined ΔRSCU values, Table 1). As shown in Table 2, the 777 genes in the top 5% average expression category (organism-wide analysis) were enriched for ribosomal protein genes and had mitochondrial and protein folding functions. We found that 14 of the 17 primary optimal codons (one per amino acid) that were previously identified using a partial transcriptome from one pooled tissue sample (embryos/ovaries [10]), were identical to those observed here, marking a strong concordance between studies and datasets (the differences herein were CAA for Gln, TTA for Leu, and AGA for Arg as optimal codons, and the presence of an optimal codon AAA for Lys, which had no optimal codon using previous embryonic/ovary data [10]). Thus, the present analysis using large-scale RNA-seq from nine divergent tissues (Additional file 1: Table S1) and using a complete annotated genome [65] support a strong preference for AT3 codons in this cricket.

**Table 1.**
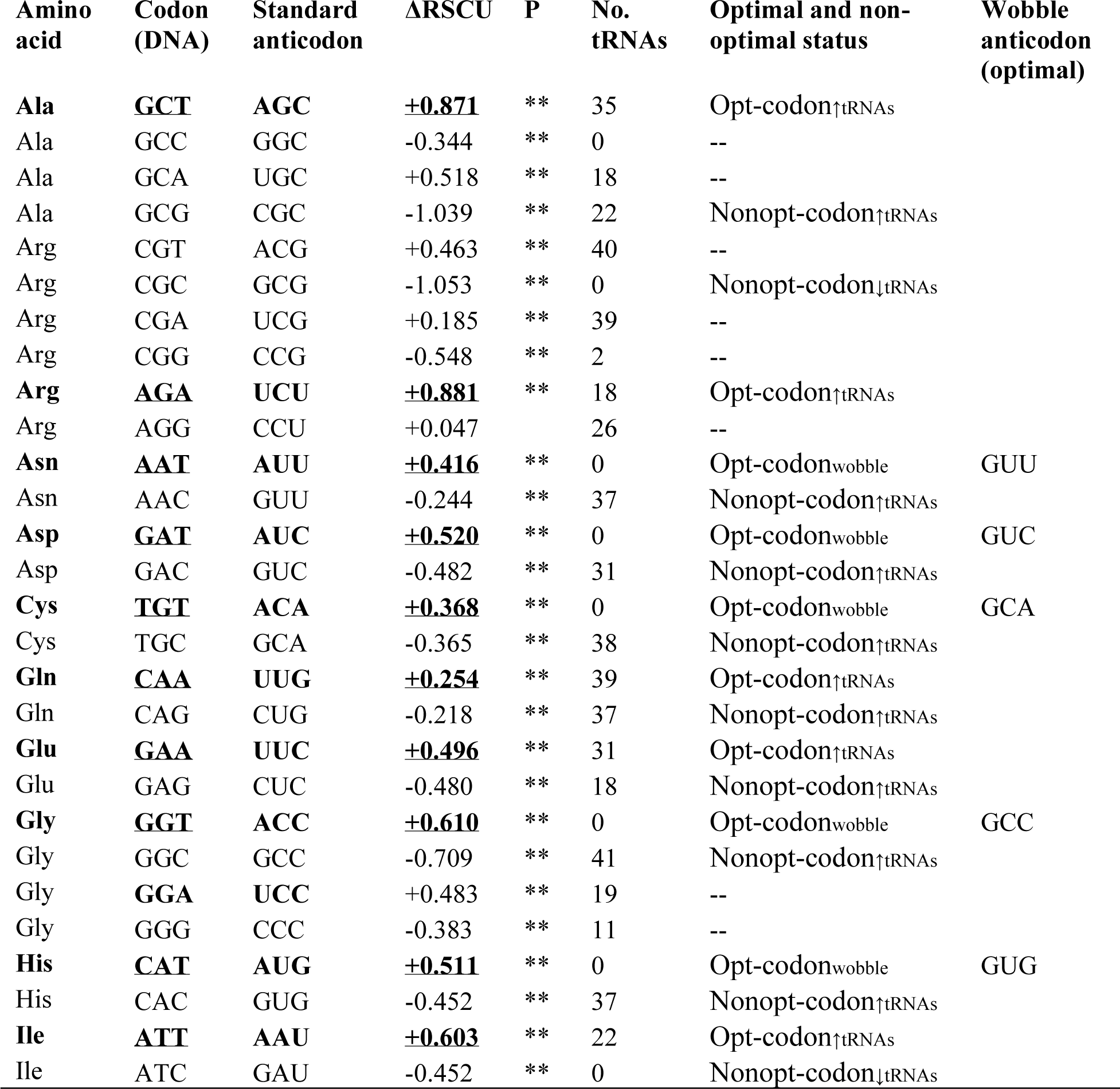
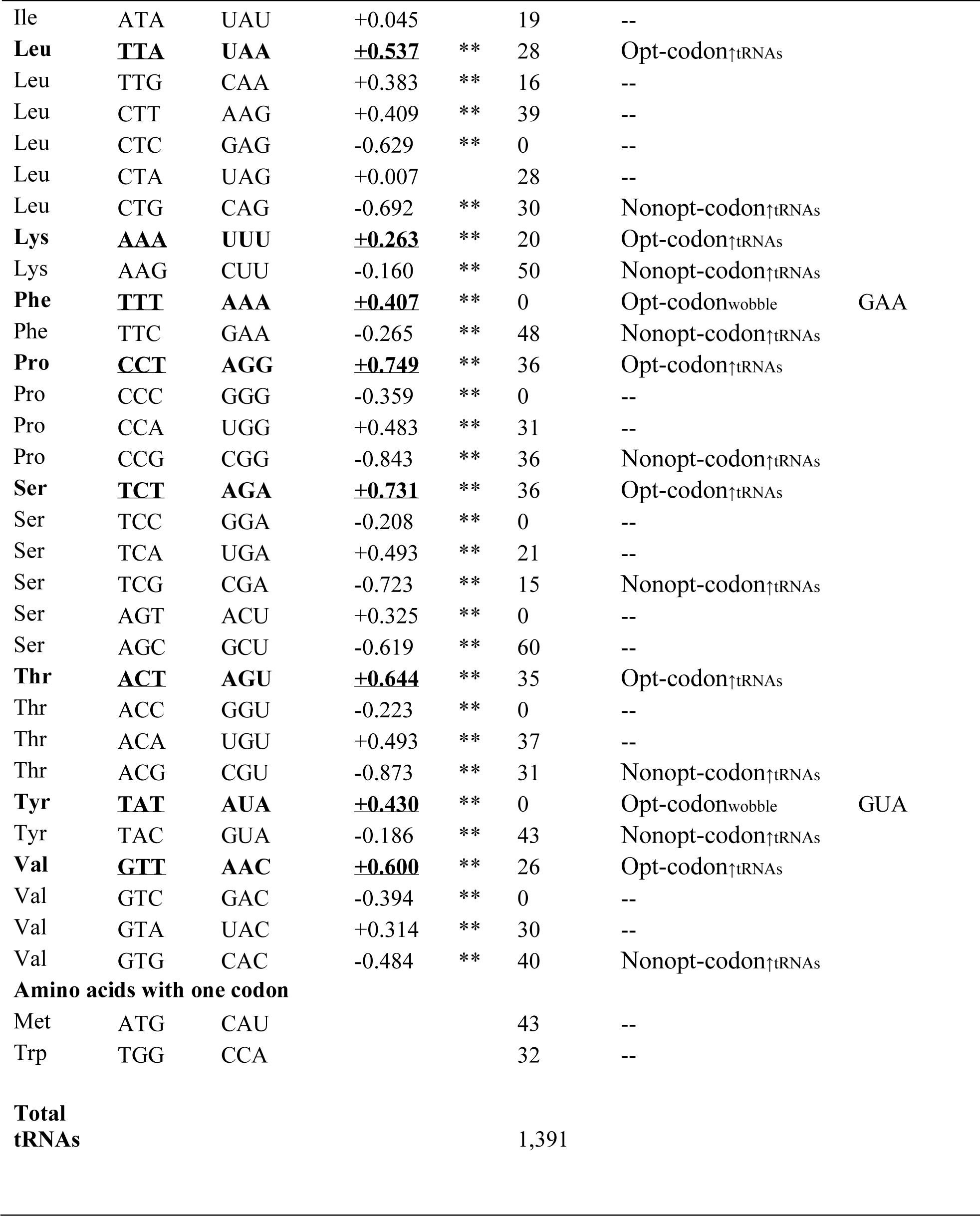
The organism-wide ΔRSCU values determined using genes with the top 5% expression level (when averaged across all nine tissues) and lowest 5% expression level (*P<0.05 **P<0.001). The number of putative tRNAs as determined using tRNA-scan and Euk filter [83] are shown. The primary optimal codon per amino acid and its ΔRSCU value are in bold and underlined. An optimal codon that has a relatively high number of tRNAs (≥18, Opt-codon_↑tRNAs_) and those with no tRNAs, and thus obligately requiring the use of wobble tRNAs (Opt-codon_wobble_), are shown, as well as the putative wobble anticodon. The primary non-optimal codons that have matching tRNA gene numbers substantially in excess of 0 (≥15, Nonopt-codon_↑tRNAs_) and those with few/no tRNAs (Nonopt-codon_↓tRNAs_) are indicated. Codons not having primary optimal or non-optimal status are indicated by “--”.

**Table 2.** Top predicted GO functional groups for organism-wide highly expressed genes (top 5% expression levels when averaged FPKM across all nine tissues). The top five clusters with the greatest enrichment (abundance) scores are shown. P-values are derived from a modified Fisher’s test, where lower values indicate greater enrichment. Data is from DAVID software [88] using those *G. bimaculatus* genes with *D. melanogaster* orthologs (BLASTX e <10^−3^ [87]).

|  |  |
| --- | --- |
| **Enrichment Score: 18.88** | P-value |
| Ribosomal protein | 7.30E-31 |
| Cytosolic ribosome | 9.00E-11 |
| **Enrichment Score: 12.49** |  |
| Mitochondrion | 3.50E-17 |
| **Enrichment Score: 8.98** |  |
| Transit peptide: Mitochondrion | 4.30E-04 |
| **Enrichment Score: 8.39** |  |
| Electron transport | 1.90E-10 |
| **Enrichment Score: 6.49** |  |
| Protein folding | 2.40E-10 |

Importantly, the expression datasets herein (Additional file 1: Table S1) allowed us to conduct an assessment of whether the identity of optimal codons varied with tissue type or sex. As certain data suggest that codon use may be influenced by the tissue in which it is maximally transcribed [16, 30], we examined those genes that exhibited maximal expression (in the top 5%) within each tissue type, that were not in the top 5% for any of the other eight remaining tissue types [16, 30], which we refer to as Top5_One-tissue_ (N values as follows, female gonad (274), male-gonad (270), female somatic reproductive system (67), male somatic reproductive system (104), female brain (24), male brain (22); female ventral nerve cord (32), male ventral nerve cord (33), and male accessory glands (162)). We found remarkable consistency among tissues, with nearly all identified optimal codons (largest positive ΔRSCU and P<0.05) ending in A3 and T3 in each tissue (Additional file 1: Table S2). For amino acids with two codons, the organism-wide optimal codon was always optimal across all nine tissues (Additional file 1: Table S2; with possible exceptions for the optimal codons AGG for Arg and CAG for Gln in the male brain; however this had P>0.1, and the N values and thus statistical power was lowest for the male brain; Additional file 1:Table S2). Nonetheless, there was some minor variation among the AT3-ending codons for amino acids with three or more synonymous codons. As an example, for the amino acid Thr, ACT was the optimal codon at the organism-wide level (Table 1) and for five tissues types (male somatic reproductive system, male brain, male ventral nerve cord, female ventral nerve cord, and male accessory glands), while the secondary organism-wide optimal codon ACA (secondary status is based on their magnitude of +ΔRSCU values) was the primary optimal codon in four other tissues (Additional file 1: Table S2). Thus, for some amino acids there is mild variation in primary and secondary status among tissues of the AT3 codons, which may reflect modest differences in the tRNA abundances among tissues [16, 32]. However, the overall patterns suggest there is remarkably high consistency in the identity of AT3 optimal codons across diverse tissues in this taxon (Additional file 1: Table S2).

While tissue-related optimal codons in multicellular organisms have only rarely been studied, the data available from fruit flies, thale cress (*Arabidopsis*), and our recent results from red flour beetles [16, 30, 32] have shown that optimal codons can vary among tissues, which suggests the existence of tissue-specific tRNA pools in those taxa [32]. The results here in *G. bimaculatus* thus differ from those in other organisms, and suggest its tRNA pools do not vary substantially with tissue or sex. Future studies using direct quantification of tRNA populations in various tissue types, which is a methodology under refinement and wherein the most effective approaches remain debated [45, 68], will help further affirm whether tRNA populations are largely similar among tissues and sex in this organism. Taken together, the results from this Top5_One-tissue_ analysis, wherein the gene set for each tissue is mutually exclusive of the top 5% expressed genes in any other tissue, suggest that high transcription in even a single tissue type or sex is enough to give rise to the optimal codons in this species. We note nonetheless that while the identity of optimal codons, and thus potentially the relative tRNA abundances, are shared among genes expressed in different tissues, the frequency of optimal codons (Fop) [22] varied among tissue types (Top5_One-tissue_), suggesting the absolute levels of tRNAs may differ among tissues (see below section “*Fop varies with tissue type and sex*”).

#### Selective pressure is a primary factor shaping optimal codons

Given that the optimal codons were highly consistent across tissues, to further investigate the potential role of selection in shaping the optimal codons we focused on the organism-wide optimal codons (Table 1). While the elevated use of the specific types of codons in highly expressed genes in Table 1 in itself provides evidence of a history of selection favoring the use of optimized codons in *G. bimaculatus* [2, 7, 9, 10, 15, 16, 66, 67, 69], the putative role of selection can be further evaluated by studying the AT (or GC) content of introns (AT-I), which are thought to largely reflect background neutral mutational pressures on genes, and thus on AT3 [16, 66, 70-74]. The *G. bimaculatus* genome contains repetitive A and T rich non-coding DNA [65], including in the introns. Nonetheless, to decipher whether any additional insights might be gained from the introns in *G. bimaculatus* we extracted the introns from the genome and found that 90.5% (N=14,071) of the 15,539 annotated genes had introns suitable for study (≥50bp after trimming). The AT-I content across all genes in this taxon had a median of 0.637, indicating a substantial background compositional nucleotide bias, and differing from the whole gene CDS (median AT for CDS across all sites=0.525). Introns (longest per gene) were nearly two-fold shorter for the most highly (top 5% organism-wide) than lowly (lowest 5%) expressed genes (1.91 fold longer in low than high expressed genes, medians were 5,183 and 2,694bp respectively, MWU-test P<0.05). We speculate that the shorter introns under high expression may comprise a mechanism to minimize transcriptional costs of abundantly produced transcripts in this cricket, as has been suggested in some other species including humans and nematodes [75], and may indicate a history of some non-neutral evolutionary pressures on the length of introns.

To further distinguish the role of mutation from selection in shaping AT3 in this cricket, we evaluated the relationship between gene expression (FPKM) and AT-I and AT3. We found that AT-I was positively correlated to gene expression level, with Spearman’s R=0.354, P<2×10^−7^ (across all 14,071 annotated genes with introns). Thus, assuming intron nucleotide content is largely selectively neutral, this may suggest a degree of expression-linked mutational-bias [76, 77] in this organism favoring AT mutations in introns of highly transcribed genes (or conversely, elevated GC mutations at low expression levels, see below in this section). However, this correlation was markedly weaker than that observed between AT3 of protein-coding genes and expression across these same genes (R=0.534, P<2×10^−7^), thus providing evidence that selection is a significant factor shaping AT3 [8].

For additional rigor in verifying the role of selection as compared to mutation in favoring AT3 codons (Table 1), genes from the top 5% and lowest 5% gene expression categories were placed into one of five narrow bins based on their AT-I content, specifically ≤0.5, >0.5-0.6, >0.6-0.7, >0.7-0.8, and >0.8. As shown in Fig. 1, for each AT-I bin, we found that AT3 of the top 5% expressed genes was statistically significantly higher than that of lowly expressed genes (MWU-tests P between 0.01 and <0.001). No differences in AT-I between highly and lowly expressed genes were observed per bin (MWU-test P>0.30 in all bins, with one exception of a minimal median AT-I difference of 0.019 for category 3, P<0.05, Fig. 1). Thus, this explicitly demonstrates that within genes that have a similar background intron nucleotide composition (that is, genes contained in one narrow bin of AT-I values), AT3 codons exhibit significantly greater use in highly transcribed than in lowly transcribed genes. This pattern further supports the interpretation that selection substantially shapes optimal codon use in *G. bimaculatus*.

**Fig. 1.**
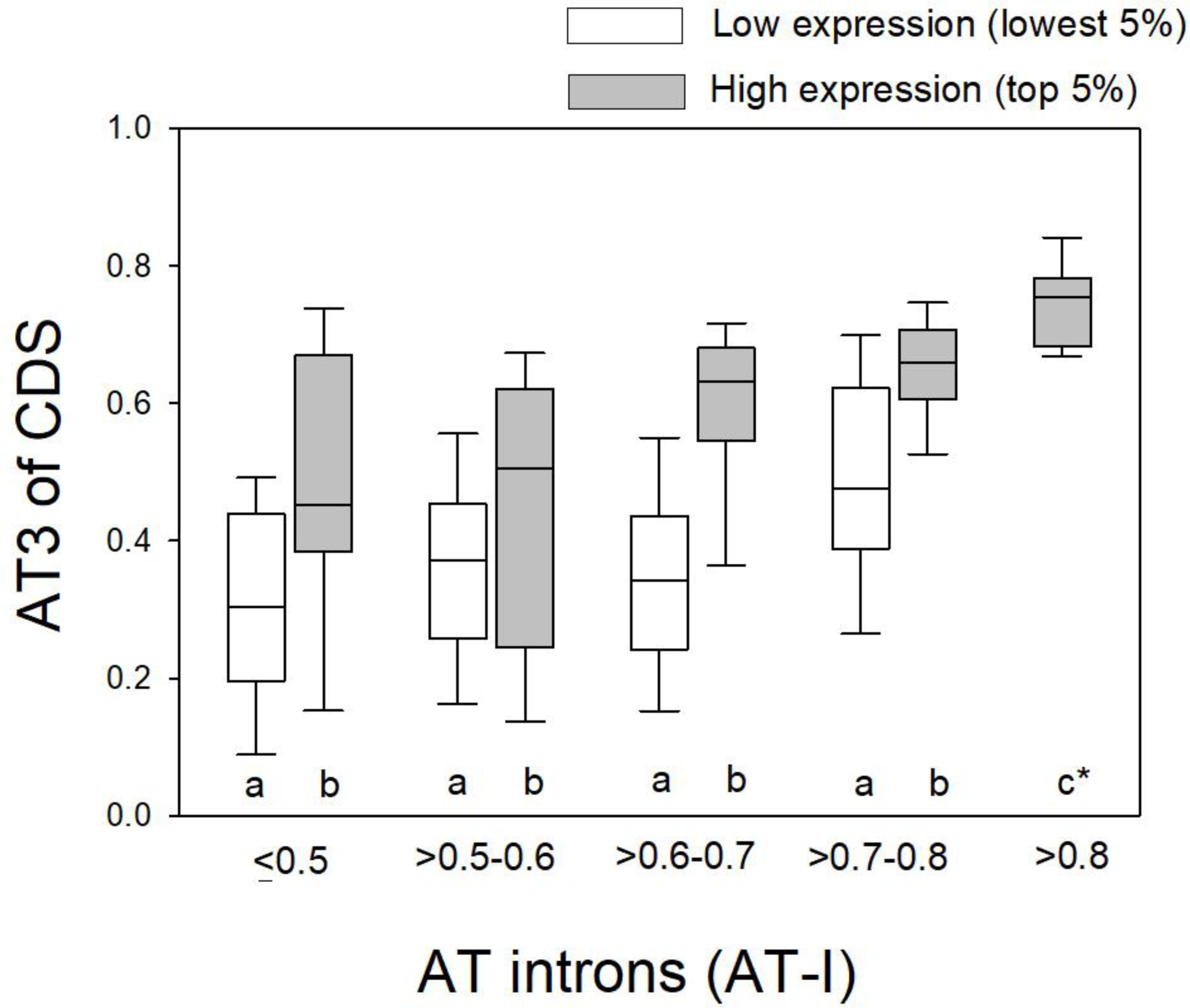
Box plots of the AT3 of codons of lowly and highly expressed genes within narrow bins of AT-I, and thus presumably having similar background mutational pressures. Genes were binned into categories with similar AT-I content to ascertain differences in AT3 attributable to non-mutational (selection) pressures in highly transcribed genes. Different letters in each pair of bars indicates P<0.05 using MWU-tests. No statistically significant differences in AT-I were observed between highly and lowly expressed genes for any bins (MWU-test P > 0.30; with the exception of a minor AT-I difference in medians of 0.019 for category 3 (0.6-0.7)). *AT3 for this bar is statistically significant from all other bars. Only one gene had AT-I >0.8 for lowly expressed genes and thus the bar for this category was excluded.

As an additional assessment, we also considered whether the lower AT3 content of lowly expressed genes (as indicated by ΔRSCU in Table 1, and in Fig. 1) could be related to biased-gene conversion, which acts to enhance GC content [74, 78], in Additional File 1: Text File S1. We conclude that while BGC may influence GC (and thus AT) content to some extent in this taxon, it is not a major factor shaping codon use of highly versus lowly expressed genes (Table 1, Additional file 1: Table S2), thereby further supporting a substantive role of selection in shaping AT3 optimal codon use patterns in Table 1 and Fig. 1.

#### Fop varies with tissue type and sex

While the types of optimal codons identified herein were largely shared among tissues (Additional file 1: Table S2), the frequency of use of these codons (Fop) varied markedly with tissue type and sex in *G. bimaculatatus*. In particular, Fop was markedly higher in Top5_One-tissue_ genes from the testes and ovaries and the male accessory glands, than in all other six tissue types (MWU-tests P<0.05, Fig. 2). Thus, this suggests that genes linked to these fundamental sexual structures and functions are prone to elevated optimal codon use that could, in principle, be due to their essential roles in reproduction and fitness, and cost-efficient translation may be particularly beneficial in the contained haploid meiotic cells [16]. Moreover, we found that the Top5_One-tissue_ genes from the female somatic reproductive system had markedly higher Fop than their male counterparts (MWU-test P<0.05, Fig. 2). We speculate that this may reflect the essential and fitness-related roles of genes involved in the insect female structures since they transport and house the male sex cells and seminal fluids after mating [79, 80], possibly making translational optimization more consequential to reproductive success for the female than male genes. In contrast, no differences in Fop were observed with respect to sex for the brain or ventral nerve cord, and the relatively low Fop values for these tissues suggest weakened selective constraint on codon use of genes as compared to the gonads and to the male accessory glands (MWU-tests P<0.05 for the latter tissues versus the former, Fig. 2). In this regard, the data show striking differences in frequency of use of the optimal codons among tissue types (Fig. 2) while the identities of optimal codons themselves are largely conserved (Additional file 1: Table S2). These patterns are consistent with a hypothesis that selection for translational optimization has been higher for genes involved in the gonads and male accessory glands, than those from the nervous system.

**Fig. 2.**
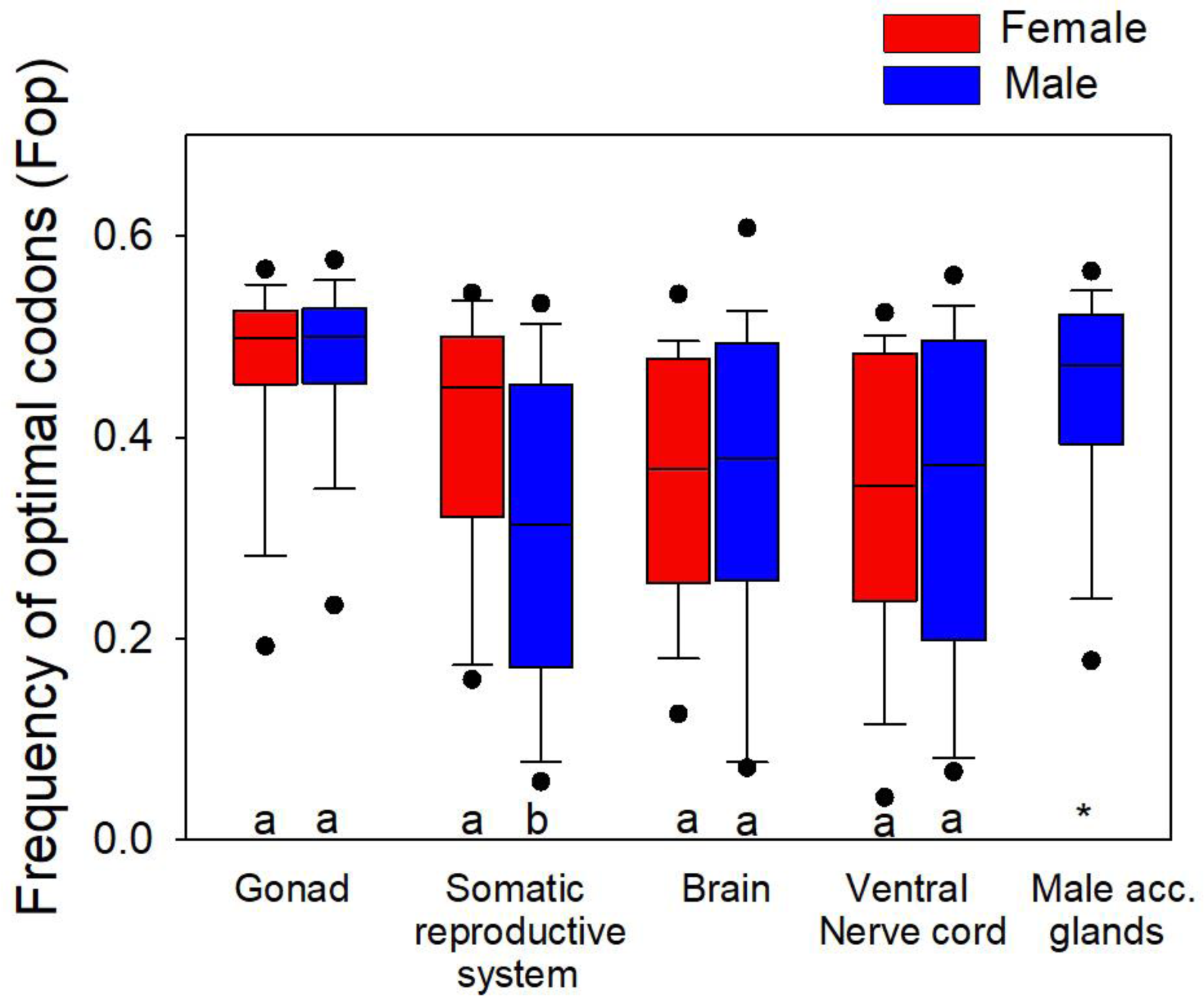
The frequency of optimal codons (Fop) for genes with expression in the top 5% in one tissue type and not in any other tissues (Top5_One-tissue_) for *G. bimaculatus*. Different letters within each pair of bars indicates a statistically significant difference (MWU-test P<0.05). Note that the gonad genes had higher Fop values than all other categories (MWU-tests P<0.05). ^*^Indicates a difference of male accessory (acc.) gland genes from all other bars.

While few comparable data on multi-tissue expression and Fop are available, and especially with respect to sex, a study of the male-female gonads and gonadectomized tissues in *D. melanogaster* indicated that codon usage bias was lower in male than female genes [31]. This pattern may be due to Hill-Robertson interference arising from adaptive evolution at linked amino acid sites in the males, dragging slightly deleterious codon mutations to fixation [31]. However, we found an opposite pattern in the mosquito *Aedes aegypti* where optimal codon use was higher in male than in female gonads [11]. Our results here, using four discrete paired male-female tissue types, suggest that the only sex-related difference in Fop for *G. bimaculatus* is for the somatic reproductive system (where male genes had lower Fop than female genes, Fig. 2). Thus, outside the somatic reproductive system, our data show that tissue type of maximal expression plays the predominant role in shaping Fop in this cricket model, rather than sex. Moreover, the low Fop observed in the brain (Fig. 2) suggests that Hill-Robertson effects may be greatest in this tissue type, a notion that is consistent with recent observations of a rapid rate of protein sequence evolution of sex-biased brain genes in this species [64]. It is worth noting that the finding that the degree of optimal codon use is particularly pronounced for genes transcribed in the gonads in Fig. 2 may suggest greater absolute (but not relative) tRNA abundances of the optimal codons in those reproductive tissues, which are essential for formation of the sex cells.

### Functional Roles of Optimal and Non-Optimal Codons Inferred by their Relationships to tRNA Gene Copies

The hypothesis of translational selection for efficient and/or accurate translation in an organism has been thought to be substantiated by associations between optimal codon use in highly expressed genes and their matching tRNA gene copy numbers in the genome [3, 5, 12, 13, 16, 20-25] In some organisms however, the correspondence between optimal codon use in highly expressed genes and the matching tRNA abundance has bean weak [21], or not observed for some (outlier) codons [81], that has been interpreted as limited support for adaptation of tRNA abundance and optimal codon use [21]. However, growing evidence suggests that there is a complex supply-demand relationship between codons and tRNAs that may affect multiple aspects of translation [42-44, 82], such that a universal connection between optimal codons and matching tRNA gene copy numbers may not always be expected [16, 42, 44]. For instance, some optimal codons may obligately require wobble tRNAs (no direct matching tRNAs) [16], which act to allow slow translation [48, 49], and thus a positive relationship between codon use in highly expressed genes and high tRNA abundance would not be expected for those codons. In turn, while non-optimal (or rare) codons may have few tRNAs, and thus act to slow translation [44], in some cases they may have numerous matching tRNAs, which could conceivably allow for translational upregulation of gene mRNAs using those codons [16, 45]. Given this context, to allow a precise interpretation of the codon-tRNA relationships in Table 1, and given some variation in terminology in the literature, we explicitly describe the codons using their ΔRSCU status and their tRNA abundances as follows: Opt-codon_↑tRNAs_ are those optimal codons (elevated use in highly expressed genes) that have relatively high tRNA gene copy numbers, Opt-codon_wobble,_ include those optimal codons obligately requiring the use of wobble tRNAs, Nonopt-codon_↓tRNAs_ are the non-optimal codons (least used in highly expressed genes) with few tRNAs, and Nonopt-codon_↑tRNAs,_ represents non-optimal codons with abundant tRNA gene copies [16].

To assess the relationships between the codon use and tRNA gene numbers for each amino acid in Table 1, we first determined the number of tRNA genes per amino acid in the *G. bimaculatus* genome using a recently updated version of the program tRNA-scan-SE (v. 2.0.5, see Methods) [83, 84]. We report 1,391 putative tRNAs for the *G. bimaculatus* genome (Table 1). To evaluate the propensity for translational selection *per se*, defined as a strong relationship between optimal codon use in highly expressed genes and tRNAs [5, 12, 18, 21], we compared the 18 primary optimal codons to the number of tRNAs per gene. We found that for 11 of 18 amino acids, the primary optimal codon had the highest or near highest matching number of tRNAs gene copies (≥18 tRNA copies) among the synonymous codons (Table 1), or Opt-codon_↑tRNAs_ status. Thus, this concurs with a model of translational selection for accurate and/or efficient translation for a majority of optimal codons in this cricket (Table 1) [5, 12, 16, 18, 21]. However, some optimal codons obligately required a wobble tRNA, or had Opt-codon_wobble_, status, which we suggest may also serve important functional roles.

#### Some optimal codons require wobble tRNAs

Seven of the 18 identified optimal codons in Table 1 had Opt-codon_wobble_ status, and had no exact matching tRNAs in the genome. These included the codons AAT (Asn), GAT (Asp), TGT (Cys), GGT (Gly), CAT (His), TTT (Phe), and TAT (Tyr) (Table 1). Thus, the elevated use of codons with Opt-codon_wobble_ status in highly transcribed genes cannot be ascribed to translational selection *per se*. We suggested in a recent report for *T. castaneum*, that optimal codons obligately using wobble tRNAs may likely be employed in highly expressed genes as a mechanism to slow translation, perhaps for protein folding purposes [16]. Indeed, experimental research in yeast, human cells, and nematodes has shown that ribosomal translocation along the mRNA is slowed by codons requiring wobble tRNAs [42, 48, 49], and thus may allow co-translational protein folding. The inefficiency of wobble interactions between codons and tRNAs, including chemically modified wobble tRNAs (e.g., adenosine to inosine, I34 in the anticodon loop [85, 86], appears to act as a mechanism to decelerate translation as compared to codons with exact tRNA matches [42, 43]. In this regard, wobble codons in highly expressed genes studied here, may serve a similar function to non-optimal codons (those that have few tRNAs, see below section), which growing studies suggest may regulate the rate, or rhythm, of translation to allow co-translational protein folding [44, 50-53]. Notably, we found the highly transcribed genes in *G. bimaculatus* were preferentially involved in protein folding as shown in Table 2, and thus this comprises a primary active process within the tissues/cells under study. In this regard, our collective results suggest a hypothesis that wobble codons in highly transcribed genes may slow translation and effectively assist in the process of protein folding.

To further study the possible roles of wobble codons, we assessed the gene ontology (GO) functions of the four codons with Opt-codon_wobble_ status that had the highest ΔRSCU values (GGT, GAT, CAT and TAT with ΔRSCU values of +0.610, +0.520, +0.511 and +0.430 respectively (Table 1)) to determine if genes using these codons tended to be involved in particular processes. For this, we examined the subset of highly expressed genes that were especially enriched for each wobble codon (had RSCU≥1.5, where a value of 1 indicates equal use of the codon per codon family, and thus ≥1.5 indicates a substantial elevation in use) in the organism-wide dataset (Table 1), and for the genes with Top5_One-tissue_ status in the gonads (Additional file 1: Table S2), which had the largest N values of any tissue type (Additional file 1: Table S2; gene ontology determined from putative orthologs to *D. melanogaster* (e<10^−3^, BLASTX [87]) and the program DAVID [88] and Flybase.org [89], see Methods). The results are shown in Additional file 1: Table S3. The functions of the organism-wide highly expressed genes with especially elevated use of the Opt-codon_wobble_ codons included ribosomal protein genes, and genes involved in mitochondrion functions (Additional file 1: Table S3), thereby specifically affirming that high use of these codons are apt to serve functions in these types of genes. For the gonads, we found that the top GO clusters for genes with high use of GAT in the ovaries (with Top5_One-tissue_ status) and of TAT in the testes (with Top5_One-tissue_ status) were involved in mitosis and cell cycle functions (Additional file 1: Table S3). Thus, this pattern for highly expressed gonadal genes in this cricket is in agreement with a prior experimental study that suggested the use of wobble codons in genes in cultured human and yeast cells might regulate the cell cycle, by controlling translation of cell-cycle genes [90]. Taken together, our results are suggestive that the use of Opt-codon_wobble_ codons in highly expressed cricket genes may act to slow translation as a means to regulate the level of cellular proteins, and to ensure proper co-translational folding, particularly affecting genes involved in the cell-cycle (Additional file 1: Table S3) and ribosomal and mitochondrial proteins (Table 2).

#### Non-optimal codons may have different functions that depend on tRNA abundance

The primary non-optimal codon per amino acid was defined as the codon with the largest negative ΔRSCU with a statistically significant P value [16]. With respect to the identified non-optimal codons, we found striking patterns with respect to tRNAs that concur with two possible functional roles, that include firstly, slowing translation, and secondly, regulating differential translation of cellular mRNAs. With respect to the former case, we found two amino acids had a primary non-optimal codon with Nonopt-codon_↓tRNAs_, status, that included CGC (Arg), ATC (Ile) (Table 1). This suggests their infrequent use in highly expressed genes may be due to the rarity or absence of matching tRNAs in the cellular tRNA pools. Moreover, these codons were not only non-optimal, and thus by definition are rare in highly transcribed genes, but their exact matching tRNAs were absent in the genome, and thus require wobble tRNAs, a combination that would in theory make them especially prone to slowing down translation. The use of non-optimal codons has been suggested to decelerate translation, which may prevent ribosomal jamming [19], and/or permit proper protein folding [44, 50, 51, 91], while, as described above, the use of codons requiring wobble tRNAs may also slow translation [42, 48, 49]. Thus, we propose the use of these two codons in genes that have Nonopt-codon_↓tRNAs_, status, and require wobble tRNAs, could play significant roles in slowing translation in highly expressed genes in *G. bimaculatus*.

Importantly however, the other non-optimal codons in Table 1 had tRNA counts markedly higher than zero (≥15 gene copies; Nonopt-codon_↑tRNAs_ status). Thus, the infrequent use of those non-optimal codons in the highly expressed genes is not likely to be due to a role in slowing translation. In fact, the use of these codons combined with high tRNA abundance suggests the potential for a high supply : demand ratio [16, 42, 45-47], a relationship that may give rise to preferential translation of any highly expressed genes that contain unusually elevated Nonopt-codon_↑tRNAs_ codons [16]. This proposed mechanism of up-translation using non-optimal (or rare) codons has been recently suggested for stress genes in yeast [45], and for highly expressed genes in the red flour beetle, wherein genes with an elevated frequency of Nonopt-codon_↑tRNAs_ status codons were linked to specific biological functions [16], suggesting their mRNAs may be preferentially translated. In this regard, the Nonopt-codon_↑tRNAs_ status codons in *G. bimaculatus* could also have significant biological roles in up-regulation of specific cellular mRNAs in this cricket model.

To further evaluable this possibility for *G. bimaculatus*, we studied as examples the Nonopt-codon_↑tRNAs_ codon GTG for Val, which had an organism-wide ΔRSCU of -0.484 and 40 tRNAs, the codon GGC for Gly with respective values of -0.709 and 41 tRNAs (note both Val and Gly are four-fold degenerate), and CTG for the six-fold degenerate Leu with a ΔRSCU of - 0.692 and 30 matching putative tRNAs (Table 1). These were chosen as examples due to their relatively high putative tRNA counts (as compared to other Nonopt-codon_↑tRNAs_ codons from amino acids with the same degeneracy level). For each of these codons, we examined those Top5_One tissue_ genes (only in the top 5% expression in one tissue type) in the gonads that had RSCU value ≥1.5, indicating enhanced use. The results are shown in Table 3. We found that genes preferentially using Nonopt-codon^↑tRNA^ codons were associated with a diverse range of functions. For example, for the ovaries, the highly expressed genes that preferentially used the Nonopt-codon_↑tRNAs_ codon GTG (for Val) included a match to *Bicaudal C* (*BicC*), which is involved in oogenesis [92]. Remarkably, this ovary gene also had elevated use of the wobble codons GGC ad CTG (Table 1). Similarly, for the ovaries, an ortholog of *santa-maria*, which has been associated with phototransduction [93] and apoptosis [94], had elevated use of each of the wobble codons GTG, GGC and CTG. The fact that both *BicC* and *santa-maria* each have high use of all three of these Nonopt-codon_↑tRNAs_ codons, which by definition have abundant matching tRNA genes, suggests their gene transcripts are preferentially translated in the ovary as compared to other transcripts in the transcript pool. For CTG (Leu), the Top5_One-tissue_ genes in the ovaries preferentially using this codon with Nonopt-codon_↑tRNAs_ status included another apoptosis gene, *apoptosis inducing factor* (*AIF*) [95], which also had elevated use of GGC for Gly, suggesting these codons may facilitate apoptosis in the female gonad cells. With respect to the testis, GTG (Val) was preferentially used in genes such as *belle*, which is involved in male germ-line stem cell development [96, 97] and *no child left behind* (*nclb*), involved in male gonad development [98], suggesting that use of this non-optimal codon may promote translation of these particular transcripts in the male gonadal mRNA pools. Enhanced use of GGC and CTG in testes genes matching *Dual-specificity tyrosine phosphorylation-regulated kinase 2* (*Dyrk2*), which is involved in apoptosis and sensory roles [99, 100], and *short spindle 3 (ssp3)*, involved in male meiosis [101] (Table 3), infers that these two codons may promote translation of apoptosis and meiotic proteins in the testes. When taken together, these patterns in *G. bimaculatus*, similar to recent findings in *T. castaneum* [16], suggest that the combination of elevated use of non-optimal codons and a high supply of tRNAs may plausibly be involved in preferential translation of the transcripts of specific genes in this system, particularly for apoptosis genes and genes with female and male gonadal functions (Table 3).

**Table 3.**
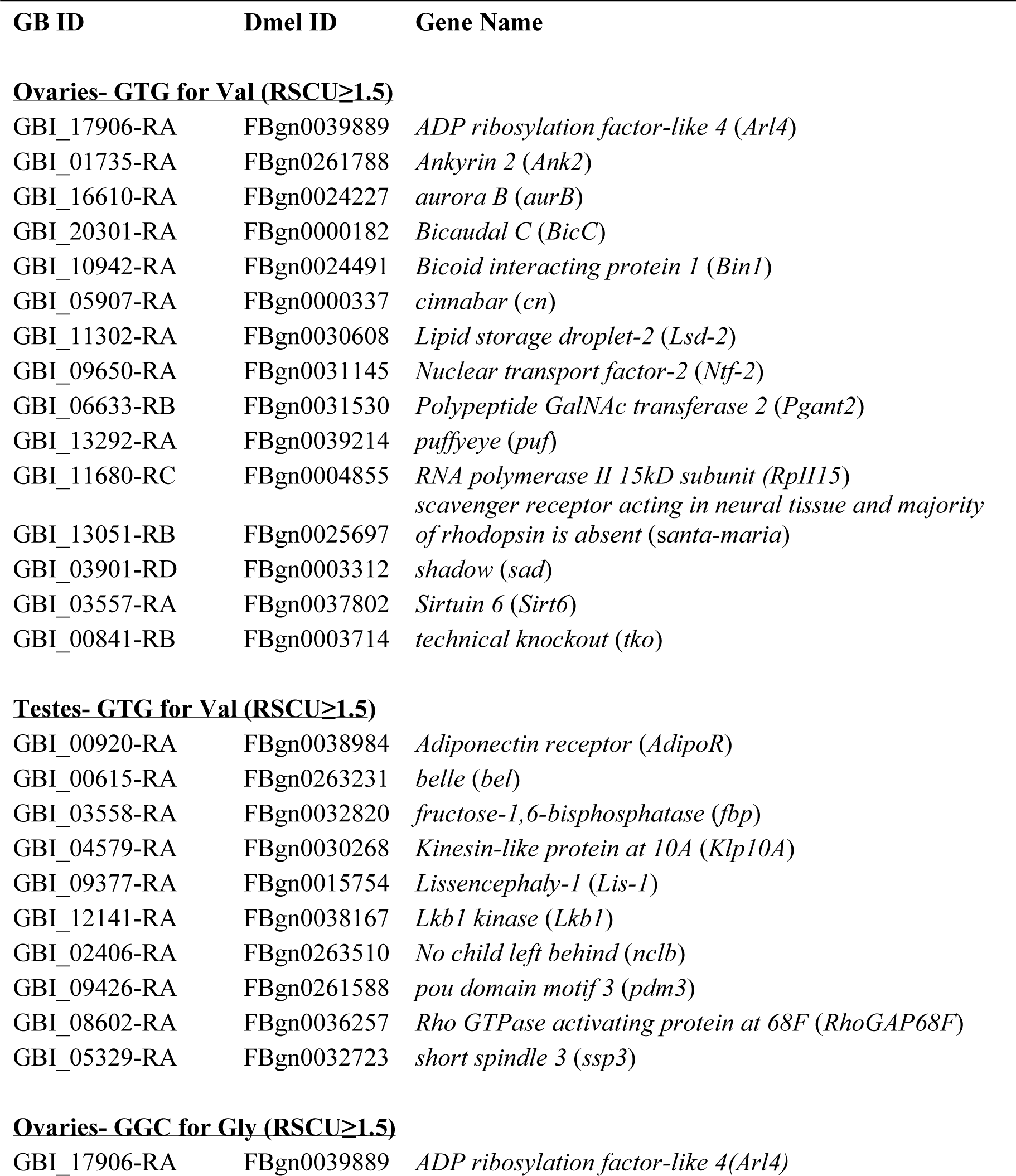
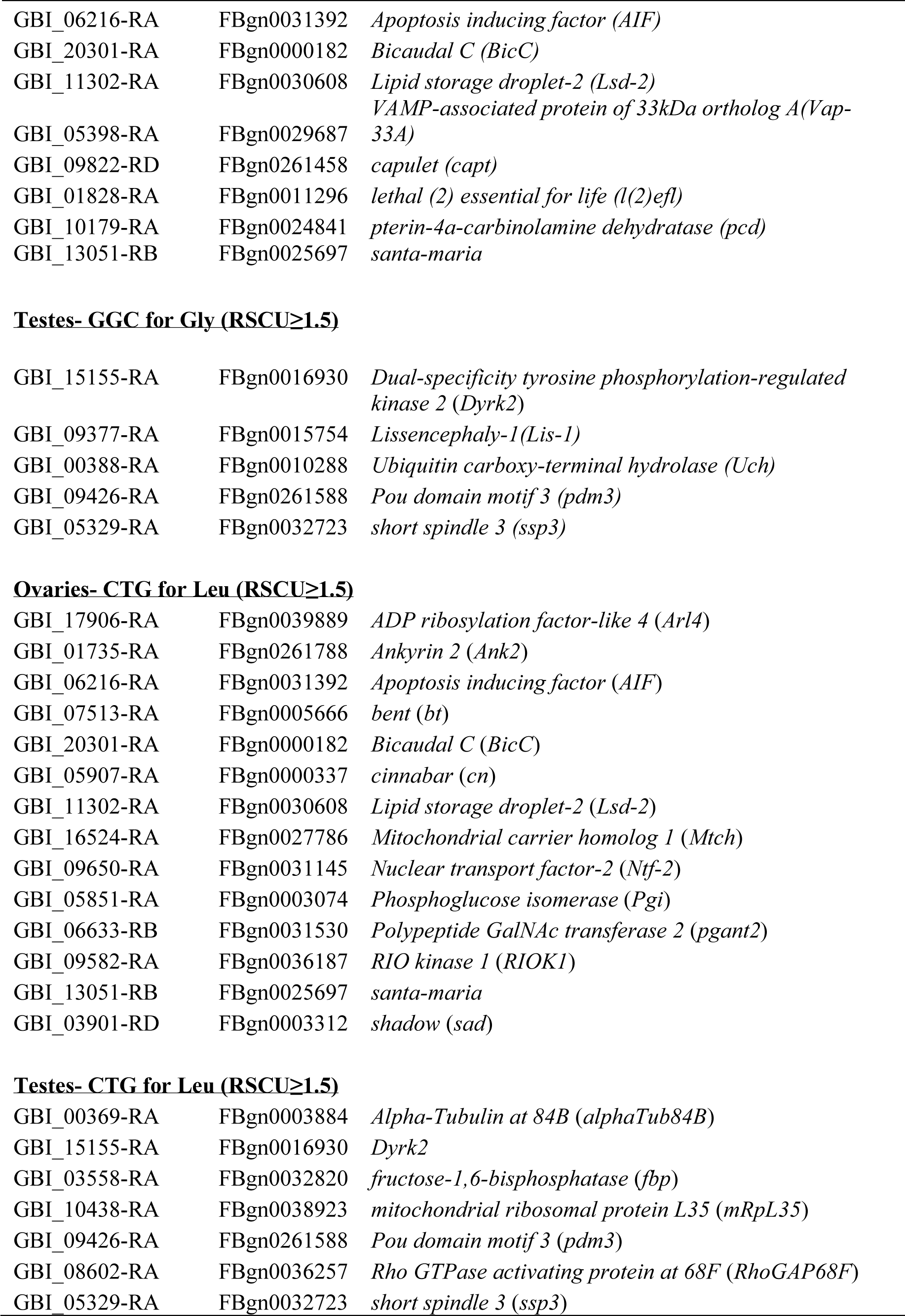
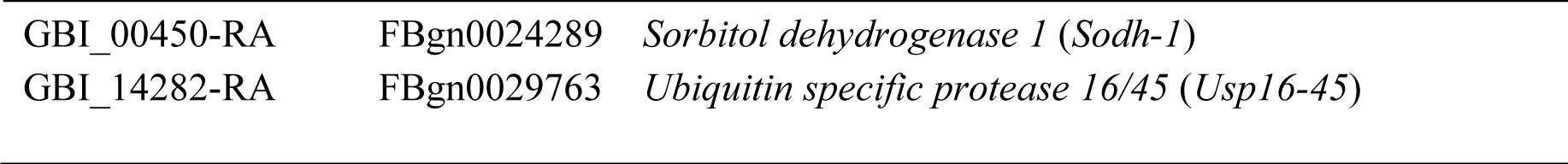
Examples of genes that exhibit the top 5% expression levels in the ovaries and top 5% expression levels in the testes (but are not in the top 5% of any other tissue type, Top5_One tissue_) in *G. bimaculatus* that have elevated use of a non-optimal codon with high tRNAs counts (Nonopt-codon_↑tRNAs_ status; elevated use in this table indicates the RSCU in a gene is ≥1.5). The codons include GTG for Val, GGC for Gly, and CTG for Leu (RSCU values ≥1.5). Genes are listed that have an identified putative *D. melanogater* (Dmel) ortholog (best match BLASTX e <10^−3^ [87] and a known gene name at FlyBase [89].

### Amino Acid Use, Biosynthesis Costs, and tRNA Gene Copies have Interdependently Evolved

Next, we asked whether amino acid use in the highly expressed genes in *G. bimaculatus* (top 5% using the organism-wide assessment) varied with their size/complexity (S/C) scores, which were developed to quantify the relative biosynthesis costs of different amino acids [56], hydropathy, or with their broad role in protein folding properties [102, 103] (Additional file 1: Table S4). As shown in Fig. 3, for highly expressed genes the amino acid usage (across all 20 amino acids) was not correlated to hydropathy (Spearman’s correlation across all 777 organism-wide highly expressed genes P>0.60) and showed no broad relationship to specific protein folding properties (ranked ANOVA P>0.05 between groups, Fig. 3BC). However, a very strong negative correlation was observed between amino acid use and S/C scores across the 20 amino acids (Spearman’s R=-0.87, P<2×10^−7^, Fig. 3A, Table 4; see also [10]). An inverse relationship between S/C score and the frequency of the 20 amino acids was also observed across all 15,539 studied *G. bimaculatus* genes irrespective of expression level (for all genes R=-0.70, P=4×10^−4^, Additional file 1: Fig. S1), but the correlation was stronger in the subset of highly expressed genes, suggesting that the connection between amino acid use and S/C scores is ameliorated with elevated transcription. Thus, these patterns both at the genome-wide level and using highly expressed genes measured across nine tissue types, indicate preferential use of low-cost amino acids in genes producing abundant mRNAs.

**Table 4.** The average amino acid use of the top 5% expressed genes (Top5_One-tissue_) in *G. bimaculatus* and 5% lowest expressed genes for the organism-wide analyses (using average expreession across all nine tissue types). **indicates P<0.05 using a two tailed t-test. The number of predicted tRNAs in the genome per amino acid are shown. SE-standard error.

| Amino acid (AA) | S/C Score | AA Freq. High exp | SE | AA Freq. Low exp | SE | Percent Diff. | P | tRNAs |
| --- | --- | --- | --- | --- | --- | --- | --- | --- |
| Gly | 1 | 6.66 | 0.21 | 8.71 | 0.13 | **-30.70** | ** | 71 |
| Ala | 4.76 | 7.32 | 0.24 | 11.54 | 0.14 | **-57.72** | ** | 75 |
| Val | 12.28 | 6.73 | 0.19 | 6.27 | 0.08 | **+6.80** | ** | 96 |
| Ile | 16.04 | 5.70 | 0.15 | 2.91 | 0.04 | **+49.01** | ** | 41 |
| Leu | 16.04 | 9.07 | 0.26 | 8.13 | 0.10 | **+10.31** | ** | 141 |
| Ser | 17.86 | 6.75 | 0.21 | 7.63 | 0.11 | **-12.94** | ** | 132 |
| Thr | 21.62 | 5.16 | 0.15 | 5.08 | 0.07 | +1.69 |  | 103 |
| Lys | 30.14 | 6.93 | 0.18 | 3.53 | 0.06 | **+49.08** | ** | 70 |
| Pro | 31.8 | 4.62 | 0.15 | 6.95 | 0.11 | **-50.40** | ** | 103 |
| Asp | 32.72 | 5.08 | 0.16 | 3.83 | 0.06 | **+24.64** | ** | 31 |
| Asn | 33.72 | 4.30 | 0.13 | 2.68 | 0.04 | **+37.70** | ** | 37 |
| Glu | 36.48 | 6.53 | 0.22 | 5.09 | 0.07 | **+22.08** | ** | 49 |
| Gln | 37.48 | 3.75 | 0.15 | 3.49 | 0.05 | **+6.92** | ** | 76 |
| Phe | 44 | 4.10 | 0.10 | 2.70 | 0.04 | **+34.20** | ** | 48 |
| Arg | 56.34 | 5.61 | 0.15 | 10.04 | 0.12 | **-78.95** | ** | 125 |
| Tyr | 57 | 3.10 | 0.08 | 1.87 | 0.05 | **+39.53** | ** | 43 |
| Cys | 57.16 | 2.08 | 0.06 | 2.51 | 0.03 | **-20.75** | ** | 38 |
| His | 58.7 | 2.24 | 0.07 | 2.53 | 0.04 | **-12.99** | ** | 37 |
| Met | 64.68 | 2.61 | 0.06 | 2.32 | 0.02 | **+10.93** | ** | 43 |
| Trp | 73 | 1.18 | 0.03 | 1.48 | 0.02 | **-25.80** | ** | 32 |

Notes: A negative correlation between S/C score and the frequency of amino acids was observed for highly and lowly expressed genes (Spearman's Ranked R=-0.87 and -0.75, P<10<sup>-7</sup>). Further, a positive correlation between the frequency of amino acids and tRNA counts was observed for highly and lowly expressed genes (Spearman's Ranked R=0.65 and 0.74, P<0.05). Percent Diff.= percent difference.

**Fig. 3.**
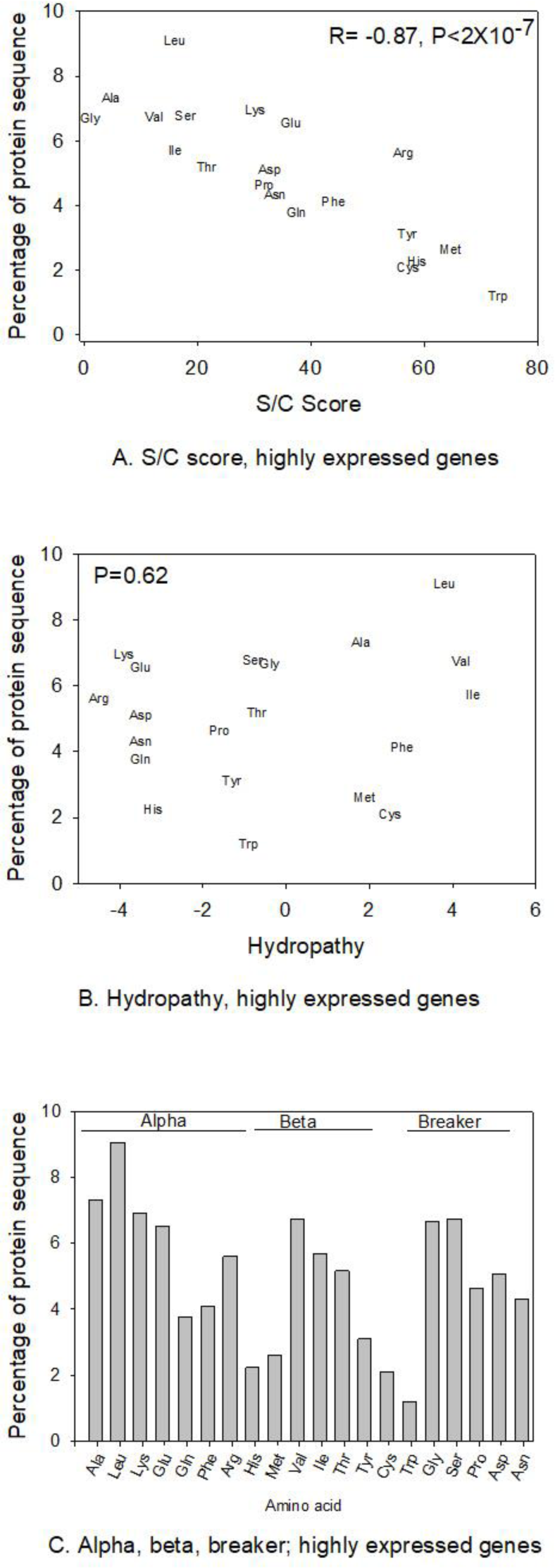
The relationship between amino acid properties and amino acid use (percent per gene, averaged across genes) in the organism-wide highly expressed genes. A) size/complexity (S/C) score; B) hydropathy, and C) folding properties. For A and B Spearman’s R and/or P values are shown, and for C no differences were detected between groups (alpha, beta, and breaker, Ranked ANOVA P>0.05).

To further decipher this relationship, we compared amino acid usage using the organism-wide highest and lowest expressed genes (top and lowest 5%, averaged across nine tissues). As shown in Table 4, we found that 19 of 20 amino acids had a statistically different frequency between the most and least transcribed genes in the genome (t-tests P<0.05), with the only exception being Thr. The amino acids with the largest increase in frequency in highly expressed genes (as compared to lowly expressed) were Ile (S/C score=16.04; with 49.0% greater use under high expression) and Lys (30.14; 49.1% greater use under high expression), suggesting that enhanced use of these amino acids with intermediate S/C scores may be more crucial to efficient translation or function of abundant transcripts, than the use of those with the lowest possible S/C scores in this taxon. We note this is consistent with an earlier analysis based on a partial transcriptome from one pooled ovary/embryo sample and without tRNA data in that study, where amino acids with intermediate S/C scores Glu, Asp, and Asn were preferred [10], that all had >22% increased use under high transcription here. This type of complex relationship between S/C score and amino acid use has also been suggested in spiders [55].

Under a null hypothesis of equal usage of each of 20 amino acids, we would assume a frequency of 5% for every amino acid per gene, with values above and below this threshold indicating favored and unfavored usage respectively. In this context, we observed that for the five highest cost amino acids (Tyr, Cys, His, Met and Trp, S/C scores of 57.00 to 73.00), the average usage was less than 5% (between 1.18 and 3.10%) in both the highly and lowly expressed genes (Table 4), indicating these biochemically costly amino acids are consistently rarely used in this taxon. Taken together, organism-wide highly expressed genes in *G. bimaculatus* exhibit a pattern of elevated use of amino acids with low S/C scores (Fig. 3A), and also exhibit elevated use of specific amino acids with intermediate S/C scores (Table 4), and very low use of the highest cost amino acids. We speculate that the pattern of favored use of some intermediate cost amino acids may be due to the roles of these amino acids in protein folding (e.g., beta and alpha folding respectively, Additional file 1: Table S4) and thus their use may ensure proper function of abundantly produced gene products.

With respect to tRNA abundances, we found that amino acid frequencies in Table 4 were positively correlated to the tRNA gene counts per amino acid (the tRNA counts included all those matching any of synonymous codons per amino acid) in *G. bimaculatus*. The correlation was observed both for the highly and for the lowly expressed genes (Spearman’s Ranked R=0.65 and 0.75, P<0.05, Table 4). Thus, this suggests the frequency of amino acid use within genes is connected to its tRNA abundance in this organism. However, despite being correlated in both groups (high and low expressed genes) in this cricket species, we suggest that the relationship is apt to be most beneficial to the organism by reducing the translational costs of genes that are highly transcribed, as these genes should presumably be most commonly translated.

We next asked whether tRNA abundance, or gene copy number, was connected to S/C scores in *G. bimaculatus*. Indeed, the 20 amino acids showed a striking tendency to be inversely connected to the total tRNA counts per amino acid in the organism-wide highly expressed genes (Spearman’s R=-0.52, P=0.02, Fig. 4). Thus, the abundance of tRNAs in the genome is directly connected to how biochemically costly an amino acid is to produce by the organism. While comparable studies of relationships between biosynthetic amino acid costs and tRNAs are uncommon, a similar negative pattern has been observed in a study from beetles [21], suggesting this phenomenon may be shared among diverse insects. Taking all our results in combination, it is evident that amino acid frequency is positively correlated to the matching tRNA gene counts (Table 4), and negatively correlated to S/C scores (Fig. 3A, Additional file 1: Fig. S1), and that tRNA gene counts per amino acid are negatively related to S/C scores (Fig. 4). In other words, genes exhibit a tendency for preferred use of low cost amino acids that have abundant tRNAs. We therefore suggest the hypothesis that all three parameters, amino acid frequency, tRNA genes in the genome, and biochemical costs, have evolved interdependently for translational optimization in *G. bimaculatus*.

**Fig. 4.**
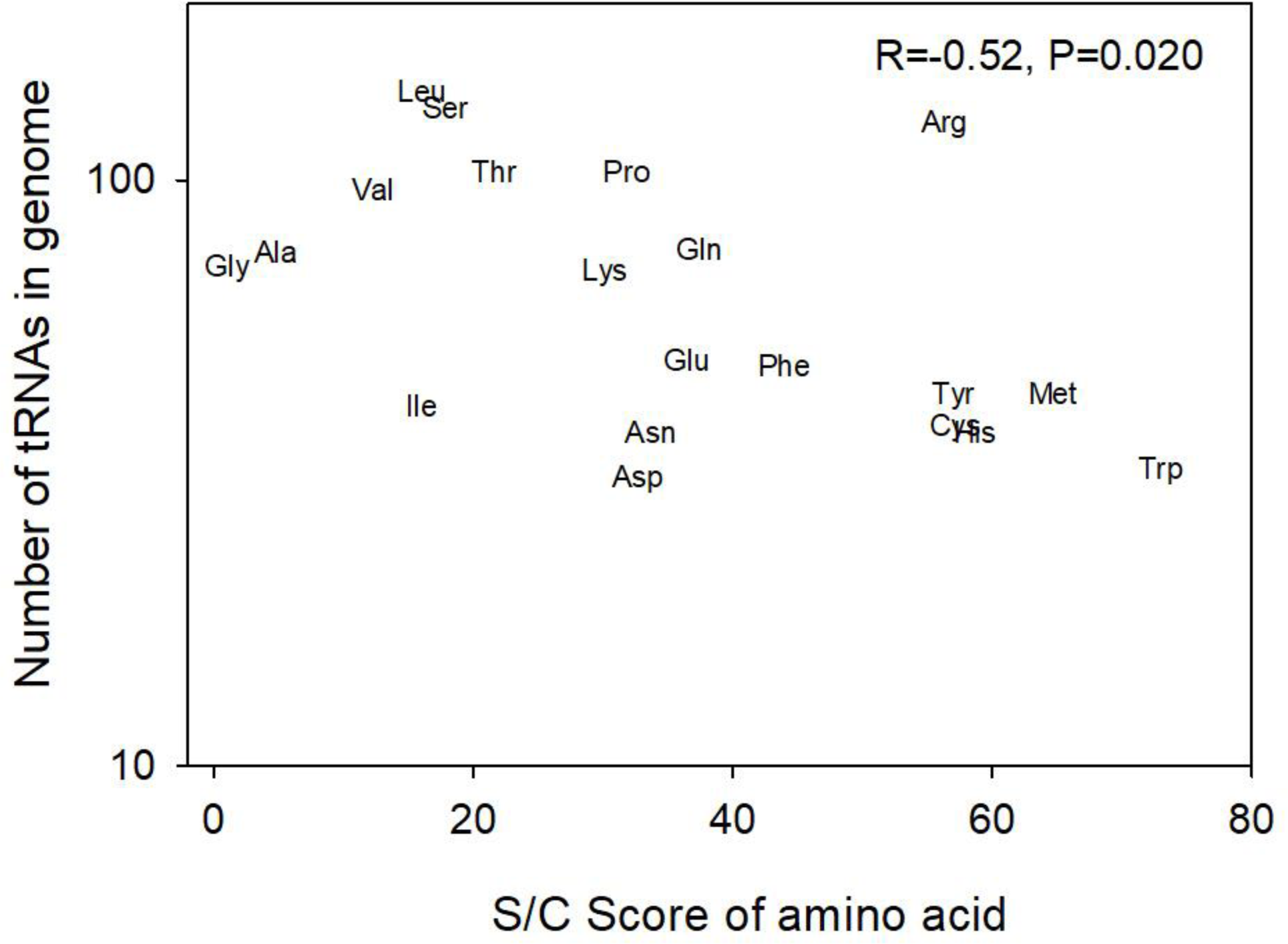
The predicted gene counts of tRNAs in the *G. bimaculatus* genome and the S/C score of each of 20 amino acids [56]. The Spearman R correlation and P value is shown.

It should be noted that while we specify herein that our tRNAs counts obtained from tRNA-scan-SE (v. 2.0.5) [83, 84] from the recently available cricket genome [65] are considered preliminary predictions in this study (see Methods, Table 1), the accuracy of this list is substantiated by the marked correlation of tRNA gene counts with S/C scores (Fig. 4) and with amino acid frequency (Table 4). In this regard, we consider the relative tRNA counts apt to provide an appropriate and accurate profile for *G. bimaculatus*.

#### Variation in amino acid use with respect to sex and tissue type

Finally, we determined whether amino acid frequency per gene varied among tissue type or sex for those genes with Top5_One-tissue_ status. The results for amino acid frequency are shown in Additional file 1: Table S5, and correlations between use for each sex per tissue type are provided in Additional file 1: Table S6. For each sex, we found strong correlations in the frequency of amino acid use (across 20 amino acids) for all paired contrasts of tissues, with Spearman R values between 0.861 and 0.98 (P<2×10^−6^). This suggests the relative amino acid use is largely consistent among highly expressed genes from all tissue types. However, the R values were weakest (R<0.9) for contrasts of the male gonad to all other tissues, suggesting a possible testis-effect on amino acid use. In terms of differences between sexes, we determined the percent difference in frequency of amino acid use between females and males for each tissue type (Additional file 1: Table S5). We found that amino acid use varied between the sexes, with between two to six amino acids per tissue type (gonad, somatic reproductive system, brain, ventral nerve cord) exhibiting statistically significant differences between sexes. As an example, for the Top5_One-tissue_ genes from the brain which had six amino acids with statistically significant differences between males and females, we found that some amino acids, namely Arg and Tyr, had in excess of 21% difference in their use between the sexes in *G. bimaculatus* (t-test P<0.05; Additional file 1: Table S5), thus revealing particularly marked variation for this tissue. In this regard, there are non-negligible differences in amino acid use between the sexes, particularly for the brain, suggesting that high expression in a particular sex may be a significant factor contributing to amino acid use.

## Conclusions

Our collective results herein strongly suggest a model whereby codon use and amino acid use have adapted to facilitate multiple functions in highly expressed genes the cricket *G. bimaculatus*. Specifically, we showed that optimal codons are largely shared across diverse tissue types and both sexes in this organism, and are likely shaped by selective pressures (Table 1, Fig. 1, Additional file 1: Table S2). Further, we revealed that a majority of optimal codons have abundant tRNA gene copies (Table 1), which is concordant with functional roles in translational optimization [12, 22]. Importantly however, we found that a substantial subset of optimal codons obligately require wobble tRNAs (Table 1), suggesting their use may have evolved as a mechanism to slow translation in highly transcribed genes, a notion that is supported by available experimental *in vivo* translation research [42, 48, 49]. These wobble codons may facilitate protein folding and/or be involved in regulation of genes such as cell-cycle genes (Additional file 1: Table S3). In turn, we demonstrated that non-optimal codons, particularly those that have few or no matching tRNAs gene copies in the genome (Table 1), may also act to slow translation, concurring with the notion that non-optimal codons may limit ribosome jamming or protein folding [19, 44, 50-53]. Crucially however, our data revealed that not all non-optimal codons are apt to have this putative function. Rather, we find that many non-optimal codons have abundant directly matching tRNA genes (Table 1), and are linked to specific types of gene ontology functions such as apoptosis and gonadal functions (Table 3). Thus, the non-optimal codons with abundant tRNAs likely provide a major organismal mechanism to promote the upregulation of specific mRNAs in the cellular mRNA pool, agreeing with a model proposed in some recent studies [16, 45]. Finally with respect to amino acid use, our data suggest a hypothesis that amino acid use, biochemical costs of amino acids [56], and tRNA gene counts in the genome (Fig. 3A, Fig. 4, Table 4) have interdependently evolved as a mechanism for translational optimization of highly expressed genes in *G. bimaculatus*.

Future research should include the direct quantification of tRNAs in different tissue types, a method that remains under development and debate [33, 45, 68, 104], to assess whether those results add support to the conclusion of similar relative tRNA abundances among amino acids across tissue types and sexes in this cricket. Moreover, further studies should be conducted of the frequencies of optimal, as well as non-optimal, codons and their relationships to tRNA abundances and gene functionalities, in a wider range of multicellular organisms. Such research will reveal whether the phenomena observed herein are shared across divergent systems.

## Materials and Methods

### Biological Samples and RNA-seq

Gene expression level was determined for all 15,539 *G. bimaculatus* protein-coding genes (CDS, longest CDS per gene) [65] that had a start codon and were >150bp. RNA-seq was obtained for four adult male and female tissue types, the gonad (testis for males, ovaries for females), somatic reproductive system, brain and ventral nerve cord and for the male accessory glands (Additional file 1: Table S1) as described previously [64]. The expression level of each *G. bimaculatus* gene was determined by mapping reads per RNA-seq dataset per tissue to the complete CDS list using Geneious Read Mapper [105], to determine FPKM per gene. FPKM was robust to mapping programs, and other common mappers including BBmap (https://jgi.doe.gov/data-and-tools/bbtools/bb-tools-user-guide/bbmap-guide/) and Bowtie2 [106] yielded similar results [64].

Optimal codons for the organism-wide analysis (averaged expression across all nine tissues) and ΔRSCU is described in the “Results and Discussion”. For each codon, using ΔRSCU=RSCU_Mean Highly Expressed CDS_-RSCU_Mean Low Expressed CDS_, t-tests were conducted between highly and lowly expressed genes to assess statistical significance. To isolate the effect of each individual tissue type, the optimal codons were determined separately for each of the nine tissues under study (males and females for each tissue type, and male accessory glands). It has been suggested that optimal codon use in a gene largely depends on the tissue type in which it is maximally transcribed [16, 30]. Accordingly, to identify optimal codons for each tissue type, we examined those genes that were in the top 5% expression in that one tissue type and not in the top 5% expression for any of the remaining eight tissues (denoted as Top5_One-tissue_) versus those with the lowest 5% expression (or all those tied with the FPKM cutoff of the lowest 5% [16]). Using these highly and lowly expressed genes per tissue, the ΔRSCU was determined as described for the organism-wide optimal codons.

The frequency of optimal codons (Fop) [4] for each gene under study was determined, using the identified optimal codons, in the program CodonW (Peden 1999). Fop was then compared for genes with high transcription in the various tissue types and two sexes in *G. bimaculatus*.

### Intron Analysis

We compared the AT (or GC) content of introns, which are thought to largely reflect the innate mutational pressures on the nucleotide content of genes [74, 107, 108], to the AT3 content (third nucleotide position) of CDS of highly and lowly expressed genes for the *G. bimaculatus* organism-wide optimal codons [16]. For this, using the genomic data for *G. bimaculatus*, we extracted the introns for all genes (with introns), and retained those >50bp after trimming of 10bp from the 5’ and 3’ ends which may contain regulatory/conserved regions [74] (and studied the longest intron per gene). For additional stringency, given that highly transcribed genes have been suggested to exhibit mutational biases (e.g., C to T) within a small number of organisms (e.g., *E. coli*, humans [76, 77]), we tested whether there was a correlation between gene expression and intron AT content in *G. bimaculatus*. To further assess the role of selection, as compared to mutation, in favoring AT3 codons (Table 1), genes from the top 5% and lowest 5% gene expression categories were placed into one of five bins based on their AT-I content as shown in Fig. 1.

### tRNA Gene Copies

The number of tRNA genes per amino acid in the *G. bimaculatus* genome was determined using the recently updated version of tRNA-scan-SE (v. 2.0.5) [83, 84]. The Eukaryotic filer called EukHighConfidenceFilter was used, which was designed to narrow the tRNA-scan output to a conservative high confidence tRNA [83] (used at default settings with the exception of ml -1). We note that since the rigor of the updated program has not been explicitly tested in insects outside *Drosophila* (P. Chan, personal communication), we consider the tRNA predictions preliminary, and focus on the relative values of tRNAs among codons and amino acids. The accuracy of the predictions, however, is strongly supported by the correlations between tRNA gene copy numbers and amino acid costs and amino acid frequency (see section “*Amino Acid Use, Biosynthesis Costs, and tRNA Gene Copies have Interdependently Evolved*”). The filter acted to reduced the absolute counts of tRNAs per amino acid to the high confidence dataset. Nonetheless, the tRNA counts with and without the filter were strongly correlated across amino acids (Spearman’s Ranked R =0.90, P<2×10^−7^), and thus relative gene counts remain intact using both measures.

### Amino Acid Use

The frequency of each of the 20 amino acids in protein-coding genes in an organism may be influenced by factors such as their size/complexity Dufton scores (which range from 1 to 73 depending on the amino acid, [56]), as well as hydropathy (where positive hydrophobicity values indicate hydrophobic nature, while negative suggest a hydrophilic amino acid [102, 103]), and/or their role in protein folding structures (alpha helices, beta sheets, or breakers used to affect bonding in helices) [103]. We thus aimed to study each of these parameters, using established values per amino acid shown in Table S4. We evaluated whether amino acid frequency in proteins of highly transcribed genes at an organism-wide level in *G. bimaculatus* (top 5% average expression across all eight male and female tissue) was correlated to S/C score [56], as well as hydropathy and protein folding characteristics [56, 102, 103]. In addition, we assessed and compared amino acid use per tissue type/sex by examining genes with Top5_One-tissue_ status per tissue type.

### Gene Ontology

For gene ontology functions, we used the gene ontology from the fly *D. melanogaster*, which comprises the most well studied insect genome to date [89]. For this, we conducted a BLAST search of the full. *G. bimaculatus* CDS list under study to *D. melanogaster* CDS list (version 6.29 [89]) using BLASTX [87], applying a cutoff of e<10^−3^. For those genes having matches within these criteria, the *D. melanogaster* gene identifiers of were then input into the program DAVID [88] for gene ontology analyses and searched in FlyBase [89].

## List of abbreviations

Top5_One-tissue_: genes with an expression level in the top 5% in one tissue type only, and not in the other eight tissues
FPKM: frequency per kilobase million
MWU-test: Mann-Whitney U-test

## Declarations

### Ethics approval and consent to participate

Not applicable.

### Consent for publication

Not applicable.

### Availability of data and material

All RNA-seq data under study are described in Additional file 1: Table S1 and are available at the Short Read Archive (SRA) under the project identifier PRJNA564136.

### Funding

This work was supported by funds from Harvard University.

#### Acknowledgements

The authors thank Dr. Guillem Ylla for providing early access to the assembled *G. bimaculatus* genome and members of the Extavour lab for discussions. The services of the Bauer core sequencing facility at Harvard University are appreciated.

## Authors’ contributions

CAW, AK and CGE designed the study. AK reared G. *bimaculatus* and sampled tissues for RNA-seq. CAW analyzed the data and wrote the manuscript with contributions by AK, NC and CGE. NC contributed to GO analysis. All authors read and approved the final manuscript.

## Additional Files

Additional File 1: The file contains the Supplementary Tables, Figures and Text which are denoted and Tables S1 to S6, Figure S1, and Text File S1.

## References

1. Plotkin JB, Kudla G: Synonymous but not the same: the causes and consequences of codon bias. Nature Reviews Genetics 2011, 12(12):32–42.

2. Whittle CA, Sun Y, Johannesson H: Evolution of synonymous codon usage in *Neurospora tetrasperma* and *Neurospora discreta*. Genome Biology and Evolution 2011, 3:332–343.

3. Percudani R, Pavesi A, Ottonello S: Transfer RNA gene redundancy and translational selection in *Saccharomyces cerevisiae*. Journal of Molecular Biology 1997, 268(268):322–330.

4. Ikemura T: Correlation between the abundance of *Escherichia coli* transfer RNAs and the occurrence of the respective codons in its protein genes: a proposal for a synonymous codon choice that is optimal for the *E. coli* translational system. Journal of Molecular Biology 1981, 151(151):389–409.

5. Akashi H: Gene expression and molecular evolution. Current Opinion in Genetics & Development 2001, 11:660–666.

6. Satapathy SS, Powdel BR, Buragohain AK, Ray SK: Discrepancy among the synonymous codons with respect to their selection as optimal codon in bacteria. DNA Research 2016, 23:441–449.

7. Ingvarsson PK: Molecular evolution of synonymous codon usage in *Populus*. BMC Evol Biol 2008, 8:307.

8. Qiu S, Bergero R, Zeng K, Charlesworth D: Patterns of codon usage bias in *Silene latifolia*. Molecular Biology and Evolution 2011, 28(28):771–780.

9. Cutter AD, Wasmuth JD, Blaxter ML: The evolution of biased codon and amino acid usage in nematode genomes. Molecular Biology and Evolution 2006, 23(23):2303–2315.

10. Whittle CA, Extavour CG: Codon and amino acid usage are shaped by selection across divergent model organisms of the Pancrustacea. G3: Genes, Genomes, Genetics 2015, 5(5):2307–2321.

11. Whittle CA, Extavour CG: Rapid Evolution of Ovarian-Biased Genes in the Yellow Fever Mosquito (*Aedes aegypti*). Genetics 2017, 206(206):2119–2137.

12. Duret L: tRNA gene number and codon usage in the *C. elegans* genome are co-adapted for optimal translation of highly expressed genes. Trends in Genetics 2000, 16(16):287–289.

13. Behura SK, Severson DW: Coadaptation of isoacceptor tRNA genes and codon usage bias for translation efficiency in *Aedes aegypti* and *Anopheles gambiae*. Insect Molecular Biology 2011, 20:177–187.

14. Whittle CA, Malik MR, Krochko JE: Gender-specific selection on codon usage in plant genomes. BMC Genomics 2007, 8:169–179.

15. Duret L, Mouchiroud D: Expression pattern and, surprisingly, gene length shape codon usage in *Caenorhabditis, Drosophila*, and *Arabidopsis*. Proc Natl Acad Sci U S A 1999, 96(96):4482–4487.

16. Whittle CA, Kulkarni A, Extavour CG: Evidence of multifaceted functions of codon usage in translation within the model beetle *Tribolium castaneum*. DNA Research 2019, 26(26):473–484.

17. Du MZ, Wei W, Qin L, Liu S, Zhang AY, Zhang Y, Zhou H, Guo FB: Co-adaption of tRNA gene copy number and amino acid usage influences translation rates in three life domains. DNA Research 2017, 24(24):623–633.

18. Sharp PM, Tuohy TM, Mosurski KR: Codon usage in yeast: cluster analysis clearly differentiates highly and lowly expressed genes. Nucleic Acids Research 1986, 14(14):5125–5143.

19. Tuller T, Carmi A, Vestsigian K, Navon S, Dorfan Y, Zaborske J, Pan T, Dahan O, Furman I, Pilpel Y: An evolutionarily conserved mechanism for controlling the efficiency of protein translation. Cell 2010, 141(141):344–354.

20. Cognat V, Deragon JM, Vinogradova E, Salinas T, Remacle C, Marechal-Drouard L: On the evolution and expression of *Chlamydomonas reinhardtii* nucleus-encoded transfer RNA genes. Genetics 2008, 179(179):113–123.

21. Williford A, Demuth JP: Gene expression levels are correlated with synonymous codon usage, amino acid composition, and gene architecture in the red flour beetle, *Tribolium castaneum*. Molecular Biology and Evolution 2012, 29(29):3755–3766.

22. Ikemura T: Codon usage and tRNA content in unicellular and multicellular organisms. Molecular Biology and Evolution 1985, 2(2):13–34.

23. Rocha EP: Codon usage bias from tRNA’s point of view: redundancy, specialization, and efficient decoding for translation optimization. Genome Research 2004, 14(14):2279–2286.

24. Moriyama EN, Powell JR: Codon usage bias and tRNA abundance in *Drosophila*. Journal of Molecular Evolution 1997, 45(45):514–523.

25. Powell JR, Moriyama EN: Evolution of codon usage bias in *Drosophila*. Proc Natl Acad Sci U S A 1997, 94(94):7784–7790.

26. Ellegren H, Parsch J: The evolution of sex-biased genes and sex-biased gene expression. Nature Reviews Genetics 2007, 8(8):689–698.

27. Ingleby FC, Flis I, Morrow EH: Sex-biased gene expression and sexual conflict throughout development. Cold Spring Harbor Perspectives in Biology 2014, 7(7):a017632.

28. Grath S, Parsch J: Sex-Biased Gene Expression. Annual Review of Genetics 2016, 50:29–44.

29. Khaitovich P, Hellmann I, Enard W, Nowick K, Leinweber M, Franz H, Weiss G, Lachmann M, Pääbo S: Parallel patterns of evolution in the genomes and transcriptomes of humans and chimpanzees. Science 2005, 309:1850–1854.

30. Camiolo S, Farina L, Porceddu A: The relation of codon bias to tissue-specific gene expression in *Arabidopsis thaliana*. Genetics 2012, 192(192):641–649.

31. Hambuch TM, Parsch J: Patterns of synonymous codon usage in *Drosophila melanogaster* genes with sex-biased expression. Genetics 2005, 170(170):1691–1700.

32. Payne BL, Alvarez-Ponce D: Codon usage differences among genes expressed in different tissues of *Drosophila melanogaster*. Genome Biology and Evolution 2019(11):1054–1065.

33. Dittmar KA, Goodenbour JM, Pan T: Tissue-specific differences in human transfer RNA expression. PLoS Genetics 2006, 2(2):e221.

34. Plotkin JB, Robins H, Levine AJ: Tissue-specific codon usage and the expression of human genes. Proc Natl Acad Sci U S A 2004, 101(101):12588–12591.

35. Liu Q: Mutational bias and translational selection shaping the codon usage pattern of tissue-specific genes in rice. PLoS One 2012, 7(7):e48295.

36. Matsumoto Y, Sakai M: Brain control of mating behavior in the male cricket *Gryllus bimaculatus* DeGeer: brain neurons responsible for inhibition of copulation actions. Journal of Insect Physiology 2000, 46(46):539–552.

37. Sakai M, Kumashiro M, Matsumoto Y, Ureshi M, Otsubo T: Reproductive Behavior and Physiology in the Cricket *Gryllus bimaculatus*. In: The Cricket as a Model Organism: Development, Regeneration and Behavior. Edited by Horch HW, Mito T, Popadic A, Ohuchi H, Noji S, vol. : Springer; 2017: 245–269.

38. Haberkern H, Hedwig B: Behavioural integration of auditory and antennal stimulation during phonotaxis in the field cricket *Gryllus bimaculatus*. Journal of Experimental Biology 2016, 219(Pt 22):3575–3586.

39. Wang B, Shao ZQ, Xu Y, Liu J, Liu Y, Hang YY, Chen JQ: Optimal codon identities in bacteria: implications from the conflicting results of two different methods. PLoS One 2011, 6(6):e22714.

40. Hershberg R, Petrov DA: Selection on codon bias. Annual Review of Genetics 2008, 42:287–299.

41. Hershberg R, Petrov DA: General rules for optimal codon choice. PLoS Genetics 2009, 5(5):e1000556.

42. Stein KC, Frydman J: The stop-and-go traffic regulating protein biogenesis: How translation kinetics controls proteostasis. Journal of Biological Chemistry 2019, 294(294):2076–2084.

43. Brule CE, Grayhack EJ: Synonymous Codons: Choose Wisely for Expression. Trends in Genetics 2017, 33(33):283–297.

44. Quax T, Claassens N, Soll D, van der Ooost J: Codon Bias as a Means to Fine-Tune Gene Expression. Molecular Cell 2015, 59:149–161.

45. Torrent M, Chalancon G, de Groot NS, Wuster A, Madan Babu M: Cells alter their tRNA abundance to selectively regulate protein synthesis during stress conditions. Sci Signal 2018, 11(11):DOI: 10.1126/scisignal.aat6409.

46. Gingold H, Dahan O, Pilpel Y: Dynamic changes in translational efficiency are deduced from codon usage of the transcriptome. Nucleic Acids Research 2012, 40(40):10053–10063.

47. Goodarzi H, Nguyen HCB, Zhang S, Dill BD, Molina H, Tavazoie SF: Modulated Expression of Specific tRNAs Drives Gene Expression and Cancer Progression. Cell 2016, 165(165):1416–1427.

48. Stadler M, Fire A: Wobble base-pairing slows in vivo translation elongation in metazoans. RNA 2011, 17(17):2063–2073.

49. Letzring DP, Dean KM, Grayhack EJ: Control of translation efficiency in yeast by codon-anticodon interactions. RNA 2010, 16(16):2516–2528.

50. Zalucki YM, Jennings MP: Experimental confirmation of a key role for non-optimal codons in protein export. Biochemical and Biophysical Research Communications 2007, 355(355):143–148.

51. Yu CH, Dang Y, Zhou Z, Wu C, Zhao F, Sachs MS, Liu Y: Codon Usage Influences the Local Rate of Translation Elongation to Regulate Co-translational Protein Folding. Molecular Cell 2015, 59(59):744–754.

52. Pechmann S, Frydman J: Evolutionary conservation of codon optimality reveals hidden signatures of cotranslational folding. Nature Structural and Molecular Biology 2013, 20(20):237–243.

53. Zhou M, Wang T, Fu J, Xiao G, Liu Y: Nonoptimal codon usage influences protein structure in intrinsically disordered regions. Molecular Microbiology 2015, 97(97):974–987.

54. Whittle CA, Extavour CG: Codon and Amino Acid Usage Are Shaped by Selection Across Divergent Model Organisms of the Pancrustacea. G3 (Bethesda) 2015, 5(5):2307–2321.

55. Whittle CA, Extavour CG: Expression-Linked Patterns of Codon Usage, Amino Acid Frequency, and Protein Length in the Basally Branching Arthropod *Parasteatoda tepidariorum*. Genome Biology and Evolution 2016, 8(8):2722–2736.

56. Dufton MJ: Genetic code synonym quotas and amino acid complexity: cutting the cost of proteins? Journal of Theoretical Biology 1997, 187(187):165–173.

57. Gaunt MW, Miles MA: An insect molecular clock dates the origin of the insects and accords with palaeontological and biogeographic landmarks. Molecular Biology and Evolution 2002, 19(19):748–761.

58. Misof B, Liu S, Meusemann K, Peters RS, Donath A, Mayer C, Frandsen PB, Ware J, Flouri T, Beutel RG et al: Phylogenomics resolves the timing and pattern of insect evolution. Science 2014, 346(346):763–767.

59. Zeng V, Ewen-Campen B, Horch HW, Roth S, Mito T, Extavour CG: Developmental gene discovery in a hemimetabolous insect: de novo assembly and annotation of a transcriptome for the cricket *Gryllus bimaculatus*. PLoS One 2013, 8(8):e61479.

60. Fisher HP, Pascual MG, Jimenez SI, Michaelson DA, Joncas CT, Quenzer ED, Christie AE, Horch HW: De novo assembly of a transcriptome for the cricket *Gryllus bimaculatus* prothoracic ganglion: An invertebrate model for investigating adult central nervous system compensatory plasticity. PLoS One 2018, 13(13):e0199070.

61. Mito T, Noji S: The Two-Spotted Cricket *Gryllus bimaculatus*: An Emerging Model for Developmental and Regeneration Studies. Cold Spring Harbor Protocols 2008, 2008:pdb emo110.

62. Donoughe S, Extavour CG: Embryonic development of the cricket *Gryllus bimaculatus*. Developmental Biology 2016, 411(411):140–156.

63. Nakamura T, Extavour CG: The transcriptional repressor Blimp-1 acts downstream of BMP signaling to generate primordial germ cells in the cricket *Gryllus bimaculatus*. Development 2016, 143(143):255–263.

64. Whittle CA, Kulkarni A, Extavour CG: Sex-biased genes expressed in the cricket brain evolve rapidly. BioRxiv 2020, www.biorxiv.org/content/10.1101/2020.07.07.192039v1

65. Ylla G, Nakamura T, Itoh T, Kajitani R, Toyoda A, Tomonari S, Bando T, Ishimaru Y, Watanabe T, Fuketa M et al: Cricket genomes: the genomes of future food. bioRxiv 2020:2020.2007.2007.191841.

66. Shields DC, Sharp PM, Higgins DG, Wright F: “Silent” sites in Drosophila genes are not neutral: evidence of selection among synonymous codons. Molecular Biology and Evolution 1988, 5(5):704–716.

67. Sharp PM, Bailes E, Grocock RJ, Peden JF, Sockett RE: Variation in the strength of selected codon usage bias among bacteria. Nucleic Acids Research 2005, 33(33):1141–1153.

68. Whittle CA, Kulkarni A, Extavour CG: Absence of a faster-X effect in beetles (*Tribolium*, Coleoptera). G3: Genes, Genomes, Genetics 2020, 10:1125–1136.

69. Cutter AD: Divergence times in *Caenorhabditis* and *Drosophila* inferred from direct estimates of the neutral mutation rate. Molecular Biology and Evolution 2008, 25(25):778–786.

70. Haddrill PR, Charlesworth B, Halligan DL, Andolfatto P: Patterns of intron sequence evolution in *Drosophila* are dependent upon length and GC content. Genome Biology 2005, 6(6):R67.

71. D’Onofrio G, Ghosh TC, Saccone S: Different functional classes of genes are characterized by different compositional properties. FEBS Letters 2007, 581(581):5819–5824.

72. Behura SK, Singh BK, Severson DW: Antagonistic relationships between intron content and codon usage bias of genes in three mosquito species: functional and evolutionary implications. Evolutionary Applications 2013, 6(6):1079–1089.

73. Zeng K, Charlesworth B: Studying patterns of recent evolution at synonymous sites and intronic sites in *Drosophila melanogaster*. Journal of Molecular Evolution 2010, 70(70):116–128.

74. Chamary JV, Hurst LD: Similar rates but different modes of sequence evolution in introns and at exonic silent sites in rodents: evidence for selectively driven codon usage. Molecular Biology and Evolution 2004, 21(21):1014–1023.

75. Castillo-Davis CI, Mekhedov SL, Hartl DL, Koonin EV, Kondrashov FA: Selection for short introns in highly expressed genes. Nature Genetics 2002, 31(31):415–418.

76. Mugal CF, von Grunberg HH, Peifer M: Transcription-induced mutational strand bias and its effect on substitution rates in human genes. Molecular Biology and Evolution 2009, 26(26):131–142.

77. Beletskii A, Bhagwat AS: Transcription-induced mutations: increase in C to T mutations in the nontranscribed strand during transcription in *Escherichia coli*. Proc Natl Acad Sci U S A 1996, 93(93):13919–13924.

78. Pouyet F, Mouchiroud D, Duret L, Semon M: Recombination, meiotic expression and human codon usage. eLife 2017, 6.

79. Degner EC, Harrington LC: A mosquito sperm’s journey from male ejaculate to egg: Mechanisms, molecules, and methods for exploration. Molecular Reproduction and Development 2016, 83(83):897–911.

80. Pascini TV, Martins GF: The insect spermatheca: an overview. Zoology 2017, 121:56–71.

81. Wright SI, Yau CB, Looseley M, Meyers BC: Effects of gene expression on molecular evolution in Arabidopsis thaliana and Arabidopsis lyrata. Molecular Biology and Evolution 2004, 21(21):1719–1726.

82. Guimaraes JC, Mittal N, Gnann A, Jedlinski D, Riba A, Buczak K, Schmidt A, Zavolan M: A rare codon-based translational program of cell proliferation. Genome Biology 2020, 21(21):44.

83. Chan PP, Lowe TM: tRNAscan-SE: Searching for tRNA Genes in Genomic Sequences. Methods in Molecular Biology 2019, 1962:1–14.

84. Lowe TM, Eddy SR: tRNAscan-SE: a program for improved detection of transfer RNA genes in genomic sequence. Nucleic Acids Research 1997, 25(25):955–964.

85. Torres AG, Pineyro D, Filonava L, Stracker TH, Batlle E, Ribas de Pouplana L: A-to-I editing on tRNAs: biochemical, biological and evolutionary implications. FEBS Lett 2014, 588(588):4279–4286.

86. Novoa EM, Pavon-Eternod M, Pan T, Ribas de Pouplana L: A role for tRNA modifications in genome structure and codon usage. Cell 2012, 149(149):202–213.

87. Altschul SF, Madden TL, Schaffer AA, Zhang J, Zhang Z, Miller W, Lipman DJ: Gapped BLAST and PSI-BLAST: a new generation of protein database search programs. Nucleic Acids Research 1997, 25(25):3389–3402.

88. Huang da W, Sherman BT, Lempicki RA: Systematic and integrative analysis of large gene lists using DAVID bioinformatics resources. Nat Protoc 2009, 4(4):44–57.

89. Gramates LS, Marygold SJ, Santos GD, Urbano JM, Antonazzo G, Matthews BB, Rey AJ, Tabone CJ, Crosby MA, Emmert DB et al: FlyBase at 25: looking to the future. Nucleic Acids Research 2017, 45:D663–D671.

90. Frenkel-Morgenstern M, Danon T, Christian T, Igarashi T, Cohen L, Hou YM, Jensen LJ: Genes adopt non-optimal codon usage to generate cell cycle-dependent oscillations in protein levels. Molecular Systems Biology 2012, 8:572.

91. Zhao F, Yu CH, Liu Y: Codon usage regulates protein structure and function by affecting translation elongation speed in *Drosophila* cells. Nucleic Acids Research 2017, 45(45):8484–8492.

92. Saffman EE, Styhler S, Rother K, Li W, Richard S, Lasko P: Premature translation of *oskar* in oocytes lacking the RNA-binding protein bicaudal-C. Molecular and Cellular Biology 1998, 18(18):4855–4862.

93. Wang T, Jiao Y, Montell C: Dissection of the pathway required for generation of vitamin A and for Drosophila phototransduction. Journal of Cell Biology 2007, 177(177):305–316.

94. Herboso L, Talamillo A, Perez C, Barrio R: Expression of the Scavenger Receptor Class B type I (SR-BI) family in Drosophila melanogaster. International Journal of Developmental Biology 2011, 55(55):603–611.

95. Stambolsky P, Weisz L, Shats I, Klein Y, Goldfinger N, Oren M, Rotter V: Regulation of AIF expression by p53. Cell Death and Differentiation 2006, 13(13):2140–2149.

96. Johnstone O, Deuring R, Bock R, Linder P, Fuller MT, Lasko P: Belle is a Drosophila DEAD-box protein required for viability and in the germ line. Developmental Biology 2005, 277(277):92–101.

97. Kotov AA, Olenkina OM, Kibanov MV, Olenina LV: RNA helicase Belle (DDX3) is essential for male germline stem cell maintenance and division in Drosophila. Biochimica et Biophysica Acta 2016, 1863(6 Pt A):1093–1105.

98. Casper AL, Baxter K, Van Doren M: no child left behind encodes a novel chromatin factor required for germline stem cell maintenance in males but not females. Development 2011, 138(138):3357–3366.

99. Luebbering N, Charlton-Perkins M, Kumar JP, Lochead PA, Rollmann SM, Cook T, Cleghon V: Drosophila Dyrk2 plays a role in the development of the visual system. PLoS One 2013, 8(8):e76775.

100. Yoshida S, Yoshida K: Multiple functions of DYRK2 in cancer and tissue development. FEBS Letters 2019, 593(593):2953–2965.

101. Wormser O, Levy Y, Bakhrat A, Bonaccorsi S, Graziadio L, Gatti M, AbuMadigham A, McKenney RJ, Okada K, El Riati S et al: Absence of SCAPER causes male infertility in humans and Drosophila by modulating microtubule dynamics during meiosis. Journal of Medical Genetics 2020.

102. Kyte J, Doolittle RF: A simple method for displaying the hydropathic character of a protein. Journal of Molecular Biology 1982, 157(157):105–132.

103. Sabbia V, Piovani R, Naya H, Rodriguez-Maseda H, Romero H, Musto H: Trends of amino acid usage in the proteins from the human genome. Journal of Biomolecular Structure and Dynamics 2007, 25(25):55–59.

104. Pang YL, Abo R, Levine SS, Dedon PC: Diverse cell stresses induce unique patterns of tRNA up- and down-regulation: tRNA-seq for quantifying changes in tRNA copy number. Nucleic Acids Research 2014, 42(42):e170.

105. Kearse M, Moir R, Wilson A, Stones-Havas S, Cheung M, Sturrock S, Buxton S, Cooper A, Markowitz S, Duran C et al: Geneious Basic: an integrated and extendable desktop software platform for the organization and analysis of sequence data. Bioinformatics 2012, 28(28):1647–1649.

106. Langdon WB: Performance of genetic programming optimised Bowtie2 on genome comparison and analytic testing (GCAT) benchmarks. BioData Mining 2015, 8(8):1.

107. Rao Y, Wu G, Wang Z, Chai X, Nie Q, Zhang X: Mutation bias is the driving force of codon usage in the *Gallus gallus* genome. DNA Research 2011, 18(18):499–512.

108. Guo X, Bao J, Fan L: Evidence of selectively driven codon usage in rice: implications for GC content evolution of Gramineae genes. FEBS Letters 2007, 581(581):1015–1021.

