## Additional File 1 for "Adaptation of codon and amino acid use for translational functions in highly expressed cricket genes"

Table. S1. The RNA-seq datasets for each of the male and female tissue types under study for *G. bimaculatus*. The number of reads (single-end) before and after trimming with BBduk (https://jgi.doe.gov/data-and-tools/bbtools) is shown. The data are available at the Short Read Archive (SRA) under the project identifier PRJNA564136 (study ID SRP220521, released upon publication). See also [1].

| | | | **No. Reads** | |
|---|---|---|---|---|
| **Sex** | **Tissues** | **Sample Name** | **Before trimming** | **After trimming** |
| Male 1 | Accessory gland | AK-28_S6.R1 | 8,519,999 | 8,455,381 |
| | Brain | AK-25_S3.R1 | 10,927,264 | 10,543,501 |
| | Somatic reproductive system | SHC-18_S14.R1 | 32,497,283 | 32,430,843 |
| | Testes | SHC-17_S13.R1 | 19,928,912 | 19,751,731 |
| | Ventral nerve cord | AK-26_S4.R1 | 11,488,521 | 11,140,299 |
| Male 2 | Accessory gland | AK-35_S13.R1 | 15,110,718 | 14,973,668 |
| | Brain | AK-32_S10.R1 | 18,039,328 | 17,850,399 |
| | Somatic reproductive system | AK-31_S9.R1 | 11,993,680 | 11,702,596 |
| | Testes | AK-30_S8.R1 | 13,672,147 | 13,529,248 |
| | Ventral nerve cord | AK-33_S11.R1 | 11,677,747 | 11,445,159 |
| Female 1 | Brain | AK-39_S17.R1 | 13,920,966 | 13,750,206 |
| | Ovary | AK-37_S15.R1 | 21,725,208 | 21,128,416 |
| | Somatic reproductive system | AK-38_S16.R1 | 13,870,827 | 13,718,497 |
| | Ventral nerve cord | AK-40_S18.R1 | 12,599,661 | 12,341,413 |
| Female 2 | Brain | AK-45_S23.R1 | 19,312,301 | 19,036,974 |
| | Ovary | AK-43_S21.R1 | 27,627,122 | 27,049,583 |
| | Somatic reproductive system | AK-44_S22.R1 | 11,688,814 | 11,539,571 |
| | Ventral nerve cord | AK-46_S24.R1 | 13,591,143 | 13,143,568 |

Table S2. The ΔRSCU for each of the nine tissues under study using genes with $Top5_{One\text{-}tissue}$ status per tissue type (versus genes with the lowest 5% expression level per tissue type). *P<0.05,**P<0.001. Note that the nongonadal tissues had fewer genes with $Top5_{One\text{-}tissue}$ expression than those with gonadal expression, particularly for the brain, and thus inherently had lower power of t-tests. However, the largest ΔRSCU per amino acid for each of the nine tissues is underlined and in bold face for all tissues irrespective of shown P value to show the tendency for high congruency among tissues. Codons reported previously as optimal codons using a pooled embryo/ovary EST dataset (Emb/Ov) are shown with an "X" [2]. N values for the $Top5_{One\text{-}tissue}$ genes are as follows: ovary (274), testis (270), female somatic reproductive system (67), male somatic reproductive system (104), female brain (24), male brain (22), female ventral nerve cord (32), male ventral nerve cord (33), and male accessory gland (162).

| Amino Acid | Codon | Organism wide | P | Ovary | P | Testis | P | Fem somatic reproductive system | P | Male somatic reproductive system | P | Female brain | P | Male brain | P | Fem ventral nerve cord | P | Male ventral nerve cord | P | Male Acc. Gland | P |
|---|---|---|---|---|---|---|---|---|---|---|---|---|---|---|---|---|---|---|---|---|---|
| Ala | GCT | **+0.871** | ** | **+0.879** | ** | **+0.914** | ** | **+0.723** | ** | **+0.249** | * | **+0.483** | * | **+0.663** | * | **+0.339** | * | **+0.376** | ** | **+0.773** | ** |
| Ala | GCC | -0.344 | ** | -0.584 | ** | -0.650 | ** | -0.375 | ** | +0.002 | | -0.154 | | -0.299 | * | -0.158 | | -0.182 | * | -0.510 | ** |
| Ala | GCA | +0.518 | ** | +0.756 | ** | +0.836 | ** | +0.416 | ** | +0.140 | | +0.342 | ** | +0.370 | * | +0.147 | * | +0.292 | ** | +0.536 | ** |
| Ala | GCG | -1.039 | ** | -1.041 | ** | -1.104 | ** | -0.750 | ** | -0.378 | ** | -0.652 | ** | -0.714 | * | -0.304 | * | -0.465 | ** | -0.839 | ** |
| Arg | CGT | +0.463 | ** | +0.387 | ** | +0.437 | ** | +0.442 | ** | +0.041 | | **+0.490** | | +0.111 | | +0.101 | | -0.025 | | +0.236 | ** |
| Arg | CGC | -1.053 | ** | -1.552 | ** | -1.658 | ** | -0.801 | ** | -0.299 | | -0.537 | | -0.548 | | -0.597 | ** | -0.780 | * | -1.295 | ** |
| Arg | CGA | +0.185 | ** | +0.183 | * | +0.364 | ** | +0.137 | * | +0.027 | | +0.431 | | +0.102 | | +0.081 | | +0.343 | * | +0.279 | * |
| Arg | CGG | -0.548 | ** | -0.520 | ** | -0.575 | ** | -0.379 | ** | -0.216 | * | -0.349 | * | -0.464 | * | -0.005 | | -0.226 | | -0.453 | ** |
| Arg | AGA | **+0.881** | ** | **+1.296** | ** | **+1.296** | ** | **+0.645** | ** | **+0.370** | * | +0.190 | | +0.392 | | **+0.361** | * | **+0.538** | * | **+1.123** | ** |
| Arg | AGG | +0.047 | | +0.203 | ** | +0.159 | ** | -0.014 | | +0.105 | | -0.197 | | **+0.436** | | -0.092 | | +0.187 | | +0.057 | |
| Asn | AAT | **+0.416** | ** | **+0.661** | ** | **+0.713** | ** | **+0.340** | * | **+0.086** | | **+0.306** | | **+0.262** | | **+0.226** | | **+0.252** | | **+0.610** | * |
| Asn | AAC | -0.244 | ** | -0.552 | ** | -0.594 | ** | -0.213 | * | +0.021 | | -0.176 | | -0.307 | | -0.096 | | -0.297 | * | -0.500 | |
| Asp | GAT | **+0.520** | ** | **+0.695** | ** | **+0.801** | ** | **+0.513** | ** | **+0.132** | | **+0.333** | | **+0.156** | | **+0.380** | * | **+0.366** | * | **+0.588** | ** |
| Asp | GAC | -0.482 | ** | -0.669 | ** | -0.761 | ** | -0.465 | ** | -0.161 | * | -0.285 | | -0.199 | | -0.333 | * | -0.374 | * | -0.563 | ** |
| Cys | TGT | **+0.368** | ** | **+0.659** | ** | **+0.698** | ** | **+0.346** | ** | **+0.155** | * | **+0.201** | * | **+0.217** | | **+0.390** | * | **+0.182** | * | **+0.504** | ** |
| Cys | TGC | -0.365 | ** | -0.594 | ** | -0.552 | ** | -0.284 | ** | -0.118 | | -0.207 | | -0.245 | | -0.236 | * | -0.142 | | -0.461 | ** |
| Gln | CAA | **+0.254** | ** | **+0.496** | ** | **+0.535** | ** | **+0.276** | ** | **+0.057** | | **+0.101** | | -0.048 | | **+0.093** | | **+0.101** | | **+0.404** | ** |
| Gln | CAG | -0.218 | ** | -0.447 | ** | -0.492 | ** | -0.224 | ** | -0.062 | | -0.048 | | **+0.098** | | -0.094 | | -0.226 | * | -0.371 | * |
| Glu | GAA | **+0.496** | ** | **+0.649** | ** | **+0.722** | ** | **+0.334** | ** | **+0.146** | * | **+0.277** | * | **+0.215** | * | **+0.209** | * | **+0.145** | | **+0.550** | ** |
| Glu | GAG | -0.480 | ** | -0.621 | ** | -0.695 | ** | -0.311 | ** | -0.142 | * | -0.337 | * | -0.284 | * | -0.183 | * | -0.238 | * | -0.527 | ** |
| Gly | GGT | **+0.610** | ** | +0.662 | ** | +0.647 | ** | **+0.485** | ** | +0.152 | | +0.350 | * | **+0.504** | * | **+0.243** | * | +0.176 | * | **+0.586** | ** |

| | | | | | | | | | | | | | | | | | | | | | |
|---|---|---|---|---|---|---|---|---|---|---|---|---|---|---|---|---|---|---|---|---|---|
| Gly | GGC | -0.709 | ** | -1.067 | ** | -1.109 | ** | -0.606 | ** | -0.139 | | -0.657 | ** | -0.431 | | -0.374 | | -0.395 | ** | -0.864 | ** |
| Gly | GGA | +0.483 | ** | **+0.714** | ** | **+0.775** | ** | +0.367 | ** | **+0.311** | * | **+0.437** | * | -0.097 | | +0.251 | | **+0.441** | * | +0.573 | * |
| Gly | GGG | -0.383 | ** | -0.320 | ** | -0.298 | ** | -0.229 | * | -0.307 | ** | -0.104 | | +0.049 | | -0.091 | | -0.190 | | -0.278 | |
| His | CAT | **+0.511** | ** | **+0.712** | ** | **+0.724** | ** | **+0.434** | ** | **+0.261** | * | **+0.192** | | **+0.205** | | **+0.160** | | **+0.405** | * | **+0.560** | ** |
| His | CAC | -0.452 | ** | -0.682 | ** | -0.646 | ** | -0.332 | ** | -0.168 | | -0.307 | | -0.342 | | -0.028 | | -0.266 | | -0.568 | ** |
| Ile | ATT | **+0.603** | ** | **+0.658** | ** | **+0.731** | ** | **+0.496** | ** | **+0.207** | | **+0.595** | * | **+0.403** | | +0.181 | | **+0.284** | | **+0.587** | ** |
| Ile | ATC | -0.452 | ** | -0.839 | ** | -0.944 | ** | -0.480 | ** | -0.081 | | -0.482 | * | -0.351 | | -0.278 | | -0.191 | | -0.709 | ** |
| Ile | ATA | +0.045 | | +0.318 | ** | +0.392 | ** | +0.062 | | -0.015 | | -0.071 | | -0.024 | | **+0.265** | | -0.100 | | +0.263 | |
| Leu | TTA | **+0.537** | ** | **+0.843** | ** | **+0.930** | ** | **+0.454** | ** | **+0.166** | * | **+0.519** | * | **+0.257** | | +0.112 | | **+0.449** | * | **+0.663** | ** |
| Leu | TTG | +0.383 | ** | +0.585 | ** | +0.553 | ** | +0.324 | | +0.102 | | +0.077 | | +0.059 | | +0.127 | | +0.130 | | +0.560 | |
| Leu | CTT | +0.409 | ** | +0.524 | ** | +0.557 | ** | +0.414 | ** | +0.041 | | +0.284 | | +0.195 | | **+0.417** | * | +0.192 | * | +0.436 | ** |
| Leu | CTC | -0.629 | ** | -0.804 | ** | -0.778 | ** | -0.492 | ** | -0.112 | | -0.358 | | -0.304 | | -0.213 | | -0.254 | | -0.625 | ** |
| Leu | CTA | +0.007 | | +0.144 | ** | +0.159 | ** | +0.086 | * | -0.008 | | +0.058 | | +0.066 | | -0.064 | | -0.065 | | +0.145 | * |
| Leu | CTG | -0.692 | ** | -1.280 | ** | -1.409 | ** | -0.778 | ** | -0.180 | | -0.576 | | -0.264 | | -0.370 | | -0.628 | * | -1.169 | ** |
| Lys | AAA | **+0.263** | ** | **+0.488** | ** | **+0.565** | ** | **+0.247** | | **+0.059** | | **+0.133** | | **+0.221** | | **+0.184** | | **+0.069** | | **+0.482** | |
| Lys | AAG | -0.160 | ** | -0.421 | ** | -0.505 | ** | -0.203 | * | +0.015 | | -0.224 | | -0.139 | | -0.173 | | -0.159 | | -0.413 | * |
| Phe | TTT | **+0.407** | ** | **+0.666** | ** | **+0.707** | ** | **+0.350** | ** | **+0.152** | ** | **+0.332** | * | **+0.309** | * | **+0.332** | * | **+0.146** | * | **+0.513** | ** |
| Phe | TTC | -0.265 | ** | -0.584 | ** | -0.614 | ** | -0.277 | ** | -0.049 | * | -0.221 | * | -0.203 | * | -0.290 | * | -0.154 | | -0.415 | ** |
| Pro | CCT | **+0.749** | ** | **+0.737** | ** | +0.828 | ** | **+0.788** | ** | **+0.279** | * | +0.351 | | **+0.364** | | **+0.452** | ** | **+0.418** | * | +0.615 | ** |
| Pro | CCC | -0.359 | ** | -0.600 | ** | -0.659 | ** | -0.504 | ** | -0.019 | | -0.292 | | -0.178 | | -0.289 | * | -0.366 | * | -0.580 | ** |
| Pro | CCA | +0.483 | ** | +0.732 | ** | **+0.873** | ** | +0.497 | ** | +0.165 | | **+0.517** | * | +0.226 | | +0.266 | * | +0.330 | * | **+0.683** | ** |
| Pro | CCG | -0.843 | ** | -0.900 | ** | -0.998 | ** | -0.727 | ** | -0.371 | ** | -0.521 | * | -0.537 | * | -0.367 | * | -0.562 | ** | -0.700 | ** |
| Ser | TCT | **+0.731** | ** | +0.691 | ** | +0.770 | ** | +0.379 | * | +0.148 | | +0.102 | | **+0.530** | | +0.141 | | +0.271 | | **+0.610** | * |
| Ser | TCC | -0.208 | ** | -0.484 | ** | -0.554 | ** | -0.326 | | +0.039 | | -0.082 | | -0.452 | * | -0.305 | * | -0.264 | | -0.479 | |
| Ser | TCA | +0.493 | ** | +0.708 | ** | **+0.855** | ** | **+0.568** | ** | **+0.350** | * | **+0.498** | * | +0.223 | | **+0.457** | * | **+0.326** | * | +0.595 | ** |
| Ser | TCG | -0.723 | ** | -0.843 | ** | -0.925 | ** | -0.551 | ** | -0.406 | ** | -0.683 | ** | -0.357 | | -0.460 | ** | -0.498 | ** | -0.696 | ** |
| Ser | AGT | +0.325 | ** | **+0.716** | ** | +0.630 | ** | +0.387 | * | +0.058 | | +0.436 | * | +0.327 | | +0.212 | | +0.424 | | +0.600 | * |
| Ser | AGC | -0.619 | ** | -0.773 | ** | -0.763 | ** | -0.443 | ** | -0.176 | | -0.258 | | -0.259 | | -0.026 | | -0.243 | | -0.619 | ** |
| Thr | ACT | **+0.644** | ** | +0.724 | ** | +0.797 | ** | +0.452 | ** | **+0.222** | * | +0.323 | * | **+0.510** | * | **+0.447** | ** | **+0.324** | | **+0.633** | ** |
| Thr | ACC | -0.223 | ** | -0.487 | ** | -0.547 | ** | -0.262 | ** | +0.050 | | -0.110 | | -0.337 | | -0.213 | | -0.106 | | -0.346 | * |
| Thr | ACA | +0.493 | ** | **+0.783** | ** | **+0.868** | ** | **+0.469** | ** | +0.163 | * | **+0.586** | ** | +0.150 | | +0.304 | * | +0.205 | * | +0.629 | ** |
| Thr | ACG | -0.873 | ** | -0.997 | ** | -1.077 | ** | -0.624 | ** | -0.439 | ** | -0.758 | * | -0.288 | | -0.501 | * | -0.498 | ** | -0.906 | ** |

| | | | | | | | | | | | | | | | | | | | | | |
|---|---|---|---|---|---|---|---|---|---|---|---|---|---|---|---|---|---|---|---|---|---|
| Tyr | TAT | **+0.430** | ** | **+0.671** | ** | **+0.668** | ** | **+0.442** | ** | **+0.203** | * | **+0.214** | | **+0.295** | | **+0.302** | * | **+0.306** | | **+0.554** | ** |
| Tyr | TAC | -0.186 | ** | -0.466 | ** | -0.477 | ** | -0.229 | * | -0.009 | | -0.156 | | -0.064 | | -0.268 | * | -0.181 | | -0.441 | * |
| Val | GTT | **+0.600** | ** | **+0.787** | ** | **+0.788** | ** | **+0.474** | ** | **+0.104** | | +0.252 | * | **+0.313** | * | +0.073 | | **+0.288** | | **+0.709** | ** |
| Val | GTC | -0.394 | ** | -0.474 | ** | -0.535 | ** | -0.361 | ** | -0.037 | | -0.136 | | +0.040 | | -0.336 | * | -0.199 | | -0.377 | ** |
| Val | GTA | +0.314 | ** | +0.435 | ** | +0.493 | ** | +0.247 | ** | +0.112 | * | **+0.302** | * | +0.255 | | **+0.196** | * | +0.138 | * | +0.347 | ** |
| Val | GTG | -0.484 | ** | -0.725 | ** | -0.741 | ** | -0.340 | ** | -0.197 | * | -0.397 | * | -0.587 | | -0.039 | | -0.329 | * | -0.661 | ** |

Table S3. Top predicted GO functional groups for organism-wide highly expressed genes (top 5% expression levels when averaged FPKM across all nine tissues) with elevated use (RSCU≥1.5) of wobble codons. Results are also shown with elevated use of the same wobble codons for genes with the top 5% expression within the ovaries and testes and not in any other tissues (Top5$_{One\text{-}tissue}$). The clusters with the greatest enrichment (abundance) scores are shown per category. P-values are derived from a modified Fisher's test, where lower values indicate greater enrichment. Data is from DAVID software [3] using those *G. bimaculatus* genes with *D. melanogaster* orthologs (BLASTX e $< 10^{-6}$ [4]).

| | | | | | |
|---|---|---|---|---|---|
| | | | **Organism wide** | | |
| **GGT Gly** | | | **GAT Asp** | | |
| Cluster 1 | Enrichment Score: 11.12 | P value | Cluster 1 | Enrichment Score: 8.77 | P value |
| | cytoplasmic translation | 2.20E-21 | | cytoplasmic translation | 3.80E-16 |
| | Ribosomal protein | 4.10E-18 | | Ribosomal protein | 1.70E-14 |
| Cluster 2 | Enrichment Score: 8.77 | | Cluster 2 | Enrichment Score: 5.56 | |
| | Mitochondrion inner membrane | 4.20E-11 | | Mitochondrion | 1.20E-09 |
| | Mitochondrion | 4.80E-10 | | Mitochondrion inner membrane | 2.00E-05 |
| Cluster 3 | Enrichment Score: 5.36 | | Cluster 3 | Enrichment Score: 4.89 | |
| | Mitochondrion | 4.80E-10 | | Mitochondrion | 1.20E-09 |
| | Transit peptide | 1.50E-05 | | Transit peptide | 2.70E-04 |
| **CAT His** | | | **TAT Tyr** | | |
| Cluster 1 | Enrichment Score: 9.8 | P value | Cluster 1 | Enrichment Score: 5.78 | P value |
| | cytoplasmic translation | 1.60E-19 | | Mitochondrion inner membrane | 6.80E-12 |
| | Ribosomal protein | 1.70E-16 | | Oxidative phosphorylation | 6.40E-08 |
| Cluster 2 | Enrichment Score: 8.61 | | Cluster 2 | Enrichment Score: 3.24 | |
| | Mitochondrion | 3.10E-11 | | Electron transport | 9.90E-05 |
| | Mitochondrion inner membrane | 1.00E-09 | | respiratory chain | 9.20E-04 |
| Cluster 3 | Enrichment Score: 6.72 | | Cluster 3 | Enrichment Score: 2.98 | |
| | Oxidative phosphorylation | 6.40E-11 | | cytoplasmic translation | 1.50E-07 |
| | Oxidoreductase | 4.00E-10 | | ribosome | 4.50E-05 |
| | | | **Top5$_{One\text{-}tissue}$ Ovaries** | | |
| **GGT Gly** | | | **GAT Asp** | | |
| Cluster 1 | Enrichment Score: 1.93 | P value | Cluster 1 | Enrichment Score: 2.35 | P value |
| | Helicase | 3.50E-04 | | eggshell chorion gene amplification | 1.30E-05 |
| | DNA/RNA helicase, DEAD/DEAH box type, N-terminal | 3.50E-03 | | Cell cycle | 5.80E-02 |
| | P-loop containing nucleoside triphosphate hydrolase | 1.10E-02 | | Cell division | 1.20E-01 |
| | ATP-binding | 5.50E-02 | Cluster 2 | Enrichment Score: 1.78 | |
| Cluster 2 | Enrichment Score: 1.38 | | | eggshell chorion gene amplification | 1.30E-05 |
| | nuclear pore | 1.10E-02 | | DNA binding | 4.00E-01 |
| | protein transporter activity | 2.70E-02 | Cluster 3 | Enrichment Score: 1.43 | |
| Cluster 3 | Enrichment Score: 1.1 | | | Protein transport | 2.90E-03 |
| | Nucleus | 8.20E-03 | | neurotransmitter secretion | 2.50E-02 |
| **CAT His** | | | **TAT Tyr** | | |
| Cluster 1 | Enrichment Score: 1.45 | P value | Cluster 1 | Enrichment Score: 1.99 | P value |
| | Zinc | 1.00E-02 | | RNA secondary structure unwinding | 2.80E-04 |
| | Metal-binding | 5.40E-02 | | RNA helicase, DEAD-box type, Q motif | 5.00E-04 |
| Cluster 2 | Enrichment Score: 1.2 | | | ATP-dependent RNA helicase activity | 1.30E-03 |
| | Protein transport | 1.90E-02 | | Nucleotide-binding | 2.60E-02 |
| | Transport | 1.10E-01 | | Hydrolase | 3.40E-01 |
| Cluster 3 | Enrichment Score: 1.04 | | Cluster 2 | Enrichment Score: 1.23 | |
| | ubiquitin-protein transferase activity | 1.40E-02 | | WD40 | 3.20E-02 |
| | Zinc-finger | 2.00E-02 | | WD40/YVTN repeat-like-containing domain | 8.80E-02 |
| | protein polyubiquitination | 8.50E-02 | Cluster 3 | Enrichment Score: 1.16 | |
| | Zinc finger, RING/FYVE/PHD-type | 2.70E-01 | | ATP-binding | 8.20E-03 |

| | | | | | |
|---|---|---|---|---|---|
| | zinc ion binding | 5.90E-01 | | Nucleotide-binding | 2.60E-02 |

**Top5$_{\text{One-tissue}}$**

**GGT Gly**

| | | |
|---|---|---|
| Cluster 1 | Enrichment Score: 1.72 | P value |
| | protein import into nucleus | 4.80E-03 |
| | Armadillo-type fold | 8.90E-03 |
| | Armadillo-like helical | 1.20E-02 |
| | protein transporter activity | 1.30E-02 |
| | cytosol | 3.70E-01 |
| Cluster 2 | Enrichment Score: 0.91 | |
| | Mitochondrion inner membrane | 4.00E-02 |
| | transmembrane region | 3.20E-01 |
| Cluster 3 | Enrichment Score: 0.67 | |
| | Transmembrane helix | 1.90E-01 |
| | Membrane | 2.00E-01 |

**GAT Asp**

| | | |
|---|---|---|
| Cluster 1 | Enrichment Score: 2.26 | P value |
| | Ubl conjugation pathway | 1.50E-03 |
| | thiol-dependent ubiquitin-specific protease activity | 3.30E-03 |
| | protein deubiquitination | 3.40E-03 |
| | Protease | 5.60E-02 |
| Cluster 2 | Enrichment Score: 1.56 | |
| | Zinc finger, RING/FYVE/PHD-type | 8.80E-03 |
| | Metal-binding | 1.90E-02 |
| Cluster 3 | Enrichment Score: 0.88 | |
| | Mitosis | 8.20E-02 |
| | Cell cycle | 2.50E-01 |

**CAT His**

| | | |
|---|---|---|
| Cluster 1 | Enrichment Score: 2 | P value |
| | Dual specificity phosphatase, subgroup, catalytic domain | 2.40E-03 |
| | protein tyrosine/serine/threonine phosphatase activity | 6.50E-03 |
| | protein dephosphorylation | 1.20E-01 |
| Cluster 2 | Enrichment Score: 1.68 | |
| | Zinc | 3.10E-03 |
| | Metal-binding | 1.70E-02 |
| | Ubl conjugation pathway | 2.90E-02 |
| Cluster 3 | Enrichment Score: 1.57 | |
| | zinc ion binding | 1.10E-02 |
| | Zinc finger, RING/FYVE/PHD-type | 1.40E-02 |
| | ubiquitin-protein transferase activity | 3.30E-02 |
| | protein ubiquitination | 5.70E-02 |

**TAT Tyr**

| | | |
|---|---|---|
| Cluster 1 | Enrichment Score: 1.63 | P value |
| | Cell cycle | 1.20E-02 |
| | Mitosis | 2.30E-02 |
| Cluster 2 | Enrichment Score: 0.95 | |
| | G-protein beta WD-40 repeat | 2.40E-02 |
| | WD40/YVTN repeat-like-containing domain | 2.30E-01 |
| Cluster 3 | Enrichment Score: 0.88 | |
| | ZnF_C2H2 | 8.20E-02 |
| | Zinc finger C2H2-type/integrase DNA-binding domain | 2.50E-01 |

Table S4. The size/complexity scores, hydropathy, and protein folding characteristics for each of the 20 amino acids. These data were used for analysis of amino acid usage [5-7].

| Amino acid | S/C score | Hydrophobic score | Folding property |
|---|---|---|---|
| Gly | 1 | -0.4 | breaker |
| Ala | 4.76 | 1.8 | alpha |
| Val | 12.28 | 4.2 | beta |
| Ile | 16.04 | 4.5 | beta |
| Leu | 16.04 | 3.8 | alpha |
| Ser | 17.86 | -0.8 | breaker |
| Thr | 21.62 | -0.7 | beta |
| Lys | 30.14 | -3.9 | alpha |
| Pro | 31.8 | -1.6 | breaker |
| Asp | 32.72 | -3.5 | breaker |
| Asn | 33.72 | -3.5 | breaker |
| Glu | 36.48 | -3.5 | alpha |
| Gln | 37.48 | -3.5 | alpha |
| Phe | 44 | 2.8 | alpha |
| Arg | 56.34 | -4.5 | alpha |
| Tyr | 57 | -1.3 | beta |
| Cys | 57.16 | 2.5 | beta |
| His | 58.7 | -3.2 | alpha |
| Met | 64.68 | 1.9 | alpha |
| Trp | 73 | -0.9 | beta |

Table S5. The average amino acid use of the Top5$_{\text{One-tissue}}$ genes in *G. bimaculatus* (frequency) for each of nine tissue types. Genes had to be in the top 5% of only one tissue type and no other tissues. Differences between male- and female-paired tissues are shown. **Indicates P<0.05 using a t-test between males and females for each tissue, *indicates P<0.1 and thus is a putative difference. Values for male accessory glands are also shown. The percent differences (Diff.) is indicated for females versus males (female demoninator). The largest three statistically signficant values per tissue are in bold.

| | | **Gonad** | | | | **Somatic reproductive system** | | | | **Brain** | | | | **Ventral nerve cord** | | | | **Accessory glands** |
|---|---|---|---|---|---|---|---|---|---|---|---|---|---|---|---|---|---|---|
| | | Frequency | | | | Frequency | | | | Frequency | | | | Frequency | | | | Frequency |
| Amino acid | S/C score | Female | Male | Diff. | P | Female | Male | Diff. | P | Female | Male | Diff. | P | Female | Male | Diff. | P | Male |
| Gly | 1 | 5.25 | 5.55 | -5.69 | * | 6.29 | 6.89 | -9.54 | | 5.79 | 7.62 | **-31.57** | * | 6.01 | 6.91 | -14.99 | | 6.75 |
| Ala | 4.76 | 6.20 | 6.29 | -1.45 | | 7.31 | 8.46 | **-15.75** | * | 7.85 | 8.68 | -10.51 | | 9.30 | 8.33 | 10.45 | | 6.91 |
| Val | 12.28 | 6.84 | 6.45 | **5.70** | ** | 6.59 | 6.61 | -0.25 | | 6.66 | 5.65 | 15.07 | | 6.83 | 6.94 | -1.62 | | 6.80 |
| Ile | 16.04 | 5.43 | 5.47 | -0.66 | | 5.01 | 4.67 | 6.73 | * | 5.14 | 5.24 | -1.91 | ** | 4.61 | 4.66 | -0.95 | | 4.23 |
| Leu | 16.04 | 9.61 | 9.37 | 2.46 | | 9.18 | 8.29 | 9.70 | | 10.12 | 8.74 | 13.61 | | 9.33 | 8.39 | 10.05 | | 9.00 |
| Ser | 17.86 | 7.94 | 7.73 | 2.68 | | 7.23 | 7.21 | 0.25 | | 7.31 | 8.63 | -18.09 | | 7.96 | 6.89 | 13.40 | | 7.47 |
| Thr | 21.62 | 5.09 | 5.13 | -0.84 | | 5.70 | 4.94 | 13.42 | | 5.58 | 4.49 | 19.46 | ** | 5.26 | 5.29 | -0.75 | | 5.01 |
| Lys | 30.14 | 6.56 | 6.73 | -2.61 | | 5.72 | 5.68 | 0.70 | | 4.91 | 5.56 | -13.19 | | 5.29 | 5.43 | -2.71 | | 4.95 |
| Pro | 31.8 | 4.51 | 4.71 | -4.43 | | 4.74 | 6.07 | **-27.88** | ** | 5.35 | 4.68 | 12.56 | | 5.65 | 6.04 | -7.00 | | 5.68 |
| Asp | 32.72 | 5.30 | 5.27 | 0.65 | | 5.83 | 4.95 | **15.14** | ** | 4.77 | 4.24 | 11.18 | | 4.96 | 4.39 | 11.50 | | 4.70 |
| Asn | 33.72 | 4.55 | 4.65 | -2.22 | | 4.35 | 4.06 | 6.71 | * | 4.03 | 3.95 | 1.86 | | 4.11 | 3.64 | 11.45 | | 3.62 |
| Glu | 36.48 | 6.82 | 7.32 | **-7.32** | ** | 6.84 | 6.20 | 9.45 | | 6.04 | 4.97 | 17.77 | | 5.97 | 6.05 | -1.25 | | 6.36 |
| Gln | 37.48 | 4.03 | 4.12 | -2.28 | | 4.12 | 3.72 | 9.85 | | 3.81 | 3.49 | 8.50 | | 4.27 | 3.66 | 14.32 | | 4.05 |
| Phe | 44 | 4.13 | 4.09 | 0.89 | | 3.54 | 3.86 | -9.05 | | 4.63 | 4.77 | -3.18 | ** | 3.81 | 3.37 | 11.51 | | 3.56 |
| Arg | 56.34 | 5.48 | 5.13 | **6.33** | * | 5.83 | 6.22 | -6.64 | ** | 5.51 | 6.68 | **-21.15** | ** | 6.80 | 7.15 | -5.09 | | 7.51 |
| Tyr | 57 | 3.09 | 2.96 | 4.10 | | 2.88 | 3.29 | -14.02 | | 3.82 | 3.00 | **21.54** | ** | 2.62 | 2.91 | -11.06 | | 2.65 |
| Cys | 57.16 | 2.44 | 2.31 | 5.12 | | 2.33 | 2.51 | -7.77 | | 1.98 | 2.72 | -37.65 | | 2.31 | 3.26 | -41.20 | | 2.44 |
| His | 58.7 | 2.53 | 2.69 | -6.03 | | 2.44 | 2.27 | 6.89 | | 2.13 | 2.70 | -27.18 | | 2.63 | 2.25 | **14.51** | * | 2.53 |
| Met | 64.68 | 2.58 | 2.55 | 1.29 | | 2.48 | 2.38 | 3.91 | | 2.68 | 2.60 | 3.06 | | 2.28 | 2.70 | **-18.63** | * | 2.37 |
| Trp | 73 | 1.23 | 1.16 | 6.00 | | 1.24 | 1.25 | -0.77 | | 1.50 | 0.94 | 36.94 | | 1.34 | 1.15 | 13.74 | | 1.28 |

Table S6. The Spearman Ranked correlations of the average amino acid frequency for genes with $Top5_{One\text{-}tissue}$ status in *G. bimaculatus*. Data were used from Table S5 determine correlations across all 20 amino acids between pairs of female and pairs of male tissues. ** Indicates P<0.001.

| **Female tissues (R-values)** | | | | | | |
|---|---|---|---|---|---|---|
| **Female tissues** | Gonad | P | SRS | P | Brain | P |
| Somatic reproductive system | 0.948 | ** | | | | |
| Brain | 0.907 | ** | 0.958 | ** | | |
| Ventral nerve cord | 0.904 | ** | 0.957 | ** | 0.956 | ** |

| **Male tissues (R-values)** | | | | | | | | |
|---|---|---|---|---|---|---|---|---|
| **Male tissues** | Gonad | P | SRS | P | Brain | P | VNC | P |
| Somatic reproductive system | 0.894 | ** | | | | | | |
| Brain | 0.884 | ** | 0.95 | ** | | | | |
| Ventral nerve cord | 0.861 | ** | 0.973 | ** | 0.938 | ** | | |
| Male Accessory glands | 0.87 | ** | 0.964 | ** | 0.925 | ** | 0.977 | ** |

Notes: SRS=somatic reproductive system, VNC=ventral nerve cord.

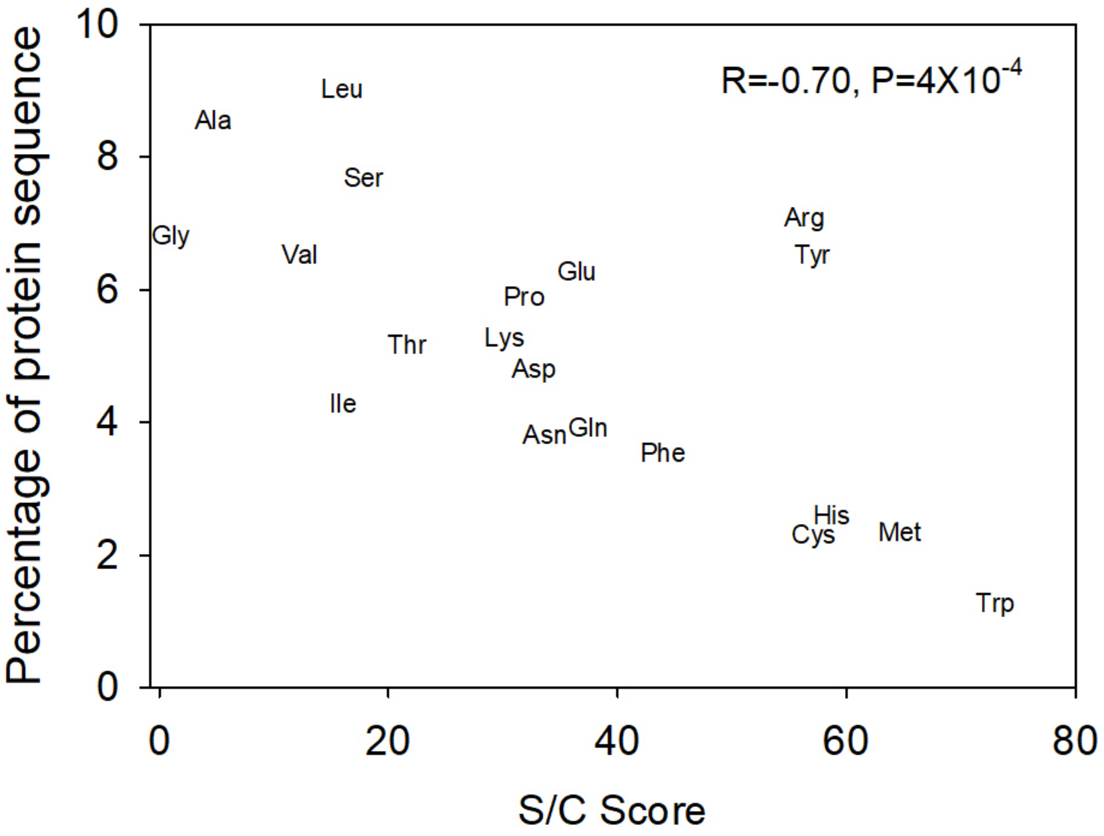


Fig. S1. The relationship between amino acid usage (percent per gene, averaged across all genes) and size/complexity (S/C) score across all 15,539 annotated genes in *G. bimaculatus*. Spearman's R and P values are shown.

**Text File S1**

Biased Gene Conversion

A possible factor that could contribute to a differences in AT3 content between low and high expressed genes, used here to define optimal codons (Table 1), is biased gene conversion (BGC) [8]. For instance, it has been reported that errors during DNA repair can lead to enhanced GC content of genes, due to favoring of GC insertions in mismatch repair of strand breaks resulting from meiotic recombination, that can ultimately enhance GC content [9-12]. In humans BGC conversion was found to be more common in lowly than highly expressed genes in the germ cells during meiosis, which was interpreted as reflecting greater crossing-over events in genes exhibiting low expression (or, inhibition of crossing-over events in genes with high expression) [13, 14]. Thus, the high AT3 (or low GC3) of organism-wide highly expressed genes observed here in *G. bimaculatus* in Table 1 could possibly result from lower BGC (as the organism-wide lowly expressed genes had <1 FPKM (or absent expression) in the male and in the female gonads containing the meiotic cells. Such genes thus may be prone to more frequent crossing over than highly expressed genes, which could cause lower and higher AT3 respectively). Further, BGC may be expected to enhance both the GC3 and GC-I content in the lowly expressed genes [13, 15], which may be consistent with a positive correlation for AT-I and AT3 to gene expression level (that infers higher GC content at lower expression, see main text). Thus, we do not exclude a role of BGC in contributing to the background nucleotide composition, and in partially shaping codon use in protein-coding genes in *G. bimaculatus*, particularly in the GC content of lowly expressed gonadal genes, as has been suggested in some mammals [13]. Crucially however, the fact that we found nearly identical AT3 optimal codons across all nine distinct tissue types, including those highly versus lowly expressed genes from male and female meiotic tissues (testis, ovary), and for all other seven studied tissues wherein meiosis does not occur (Additional file 1: Table S2), suggests BGC is not the primary factor shaping optimal codon use (or, in other words, not causing the AT3 differences between high and low expression classes). Rather, optimal codon use in highly expressed genes is likely substantially shaped by selection, with a comparatively minor role of mutational bias (Fig. 1).

**References**


1. Whittle CA, Kulkarni A, Extavour CG: **Sex-biased genes expressed in the cricket brain evolve rapidly**. *BioRxiv* 2020, www.biorxiv.org/content/10.1101/2020.07.07.192039v1
2. Whittle CA, Extavour CG: **Codon and amino acid usage are shaped by selection across divergent model organisms of the Pancrustacea**. *G3: Genes, Genomes, Genetics* 2015, **5**(11):2307-2321.
3. Huang da W, Sherman BT, Lempicki RA: **Systematic and integrative analysis of large gene lists using DAVID bioinformatics resources**. *Nat Protoc* 2009, **4**(1):44-57.
4. Altschul SF, Madden TL, Schaffer AA, Zhang J, Zhang Z, Miller W, Lipman DJ: **Gapped BLAST and PSI-BLAST: a new generation of protein database search programs**. *Nucleic Acids Research* 1997, **25**(17):3389-3402.
5. Sabbia V, Piovani R, Naya H, Rodriguez-Maseda H, Romero H, Musto H: **Trends of amino acid usage in the proteins from the human genome**. *Journal of Biomolecular Structure and Dynamics* 2007, **25**(1):55-59.
6. Kyte J, Doolittle RF: **A simple method for displaying the hydropathic character of a protein**. *Journal of Molecular Biology* 1982, **157**(1):105-132.
7. Dufton MJ: **Genetic code synonym quotas and amino acid complexity: cutting the cost of proteins?** *Journal of Theoretical Biology* 1997, **187**(2):165-173.
8. Galtier N, Roux C, Rousselle M, Romiguier J, Figuet E, Glemin S, Bierne N, Duret L: **Codon Usage Bias in Animals: Disentangling the Effects of Natural Selection, Effective Population Size, and GC-Biased Gene Conversion**. *Molecular Biology and Evolution* 2018, **35**(5):1092-1103.
9. Haddrill PR, Charlesworth B, Halligan DL, Andolfatto P: **Patterns of intron sequence evolution in *Drosophila* are dependent upon length and GC content**. *Genome Biology* 2005, **6**(8):R67.
10. de Proce SM, Zeng K, Betancourt AJ, Charlesworth B: **Selection on codon usage and base composition in *Drosophila americana***. *Biology Letters* 2012, **8**(1):82-85.
11. Zeng K, Charlesworth B: **Studying patterns of recent evolution at synonymous sites and intronic sites in *Drosophila melanogaster***. *Journal of Molecular Evolution* 2010, **70**(1):116-128.
12. Marais G: **Biased gene conversion: implications for genome and sex evolution**. *Trends in Genetics* 2003, **19**(6):330-338.
13. Pouyet F, Mouchiroud D, Duret L, Semon M: **Recombination, meiotic expression and human codon usage**. *eLife* 2017, **6**.
14. McVicker G, Green P: **Genomic signatures of germline gene expression**. *Genome Research* 2010, **20**(11):1503-1511.

15. Chamary JV, Hurst LD: **Similar rates but different modes of sequence evolution in introns and at exonic silent sites in rodents: evidence for selectively driven codon usage**. *Molecular Biology and Evolution* 2004, **21**(6):1014-1023.
